# DeltaQ: Value-Guided Hebbian Learning in Spiking Neuronal Networks for Multi-Goal Navigation

**DOI:** 10.64898/2026.06.12.731882

**Authors:** Christopher Earl, Gozde Unal, Hananel Hazan, Samuel A Neymotin

## Abstract

Animals must often navigate environments where feedback about progress toward a goal is sparse or delayed, requiring internal representations of space and memory of prior experience. The hippocampal-entorhinal system is believed to support this capability through distributed spatial representations that guide goal-directed behavior. However, many computational models of these circuits focus primarily on reproducing neural dynamics rather than demonstrating how such representations support learning on navigation tasks. We present a biologically inspired spiking neuronal network (SNN) model that combines grid-cell-derived spatial representations, ΔQ-modulated Hebbian plasticity, and context-dependent modulation to support navigation under sparse reward conditions. Grid Cell populations generate distributed spatial codes that are transformed by an Association Cell population into more spatially selective internal representations. Learning is driven by changes in Q-values (ΔQ) computed from a goal-conditioned Q-table, allowing local synaptic plasticity to incorporate information about long-term navigation outcomes. For environments containing multiple navigation objectives, a Context Cell population provides task-dependent modulation that enables a shared network architecture to support distinct navigation policies. Across two complementary maze environments, the model demonstrates three core capabilities: generation of distinct spatial representations, learning of efficient navigation policies under sparse and delayed reward, and support for multiple navigation objectives within a shared environment. The results further show that contextual modulation introduces subtle task-dependent variations into a largely shared population representation, allowing identical spatial locations to support different navigation behaviors. These findings demonstrate that biologically inspired spatial representations, value-guided plasticity, and contextual modulation can jointly support flexible navigation in spiking neuronal networks, providing a bridge between mechanistic neural circuit models and functional reinforcement learning.

## Introduction

Biological organisms can learn efficient navigation strategies in environments that provide sparse and delayed feedback. Understanding how neural circuits support such learning remains a central challenge in computational neuroscience. The hippocampal (HPC) and entorhinal cortex (EC) are known to encode spatial information through structured population activity, yet the circuit mechanisms by which these representations support goal-directed behavior are not fully understood (Moser et al., 2008; O’Keefe & Dostrovsky, 1971).

Spiking neuronal network (SNN) models using reward-modulated spike-timing-dependent plasticity (STDP/RL) have demonstrated the ability to learn a variety of sensorimotor tasks(Sanda et al., 2017; Skorheim et al., 2014), including control of simulated limbs and robotic actuators (Chadderdon et al., 2012; Florian, 2007; Neymotin et al., 2013), as well as decision-making tasks such as video game play (Anwar et al., 2022; Haşegan et al., 2022; Patel et al., 2019). In many of these settings, reward signals vary continuously with behavioral progress, providing frequent feedback that can reinforce beneficial activity patterns(Doya, 2000; Frémaux et al., 2013). Under such conditions, reward-modulated local plasticity can effectively associate neural activity with desirable outcomes.

Navigation tasks such as maze learning present a more challenging learning regime. In these environments, most actions produce little or no feedback, and reward is often obtained only after extended sequences of decisions. Consequently, learning requires associating local neural activity with delayed outcomes while simultaneously maintaining information about previously visited locations. Sparse feedback makes it difficult for conventional reward-modulated plasticity mechanisms to assign credit to earlier neural activity that contributed to successful navigation.

Reinforcement learning algorithms such as Q-learning (QL) address this challenge by estimating expected future reward for state-action pairs (Connell & Mahadevan, 1999; Montague et al., 1996). Recent work has demonstrated that Q-like learning dynamics can be implemented within neural systems, suggesting potential links between temporal-difference learning and biologically plausible plasticity mechanisms (Nejime et al., 2026; Shin et al., 2026). In the present work, Q-learning is used as an external process to compute changes in value (ΔQ) that modulate local Hebbian plasticity.

Rather than directly embedding Q-values within neural firing rates, ΔQ acts as a teaching signal that links local synaptic updates to long-term navigation outcomes while preserving locality of plasticity mechanisms.

To support navigation learning, the model incorporates neuromimetic spatial representations inspired by population coding in the EC and HPC. Distributed grid-cell-derived activity is transformed into more spatially selective internal representations that can be associated with navigation decisions through ΔQ-modulated Hebbian plasticity. Together, these components address three core challenges of navigation: (1) generating distinct representations for different spatial locations, (2) learning effective action sequences under sparse and delayed reward conditions, and (3) supporting multiple navigation objectives within a shared environment.

To evaluate these challenges, we consider two complementary maze environments that differ in structure and task demands (**Fig. 1**). Both environments contain obstacles that restrict movement and require multi-step planning to reach goal locations. Maze Type 1 focuses on learning efficient navigation toward a common goal from multiple starting locations, emphasizing spatial state discrimination and long-horizon learning. Maze Type 2 introduces multiple start-goal pairs within a shared spatial layout, requiring the same network to support multiple navigation objectives simultaneously. To address this challenge, the model incorporates a Context Cell population that provides task-dependent modulation of internal network activity.

**Figure 1:**
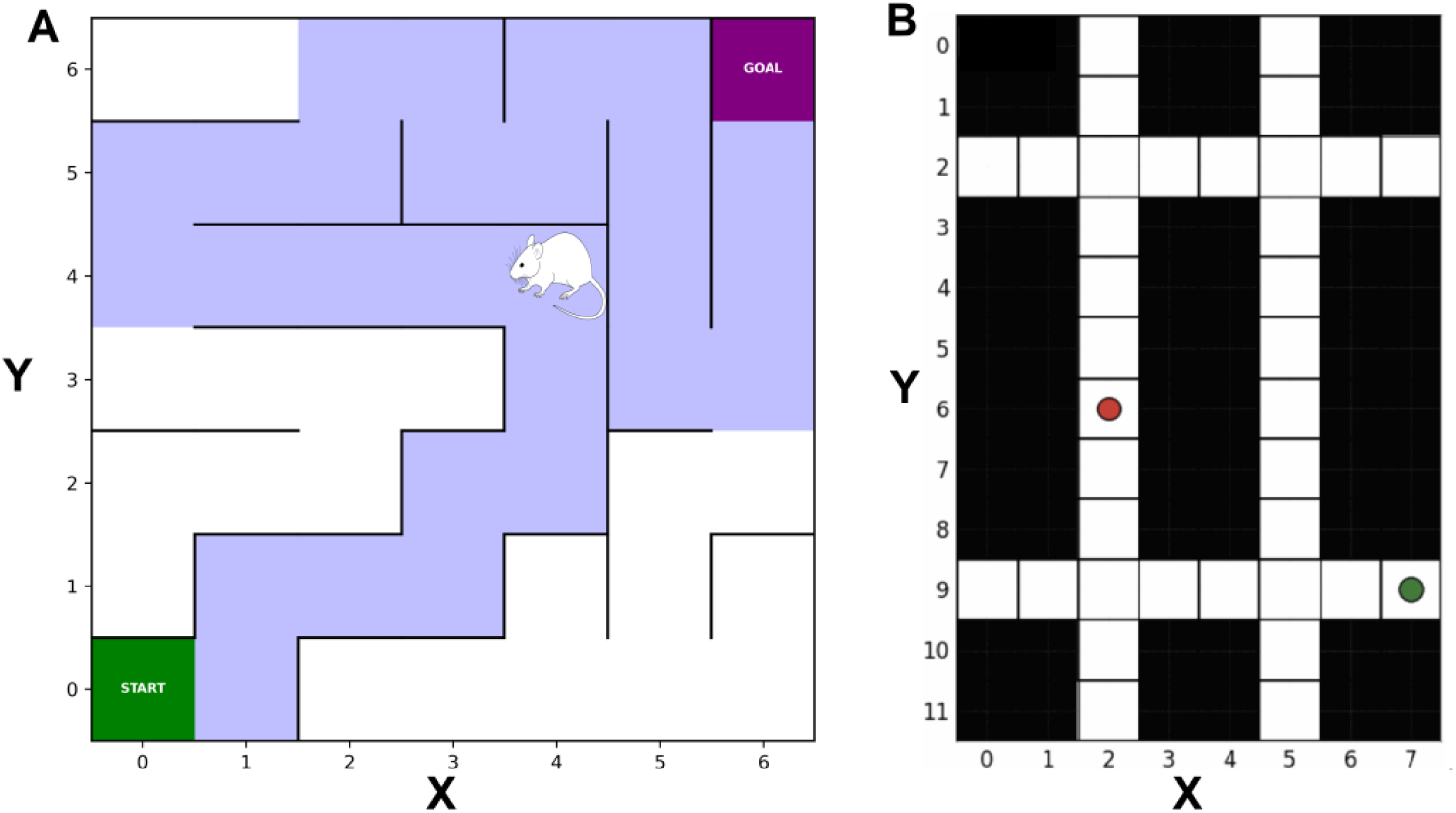
Simulated maze environments. A) 7x7 Maze Type 1 with the model-controlled agent (mouse) at coordinate position (4, 4). The maze is divided into square tiles, one for each coordinate location. The model interacts with the maze by simulating GC firing behaviors relative to its position in the maze, and outputting one of four action choices, Up, Down, Left, or Right. There are three kinds of environmental feedback: negative feedback (-1) if the agent runs into a wall, positive feedback (+1) if the agent intercepts the goal, null feedback (0) if the agent moves into an open tile. Most moves will result in null or negative feedback, hence why we classify it as a sparse feedback environment. **B)** 12x8 **Maze Type 2**, where the model-controlled agent must learn to reach the target (green circle) from the starting location (red circle). The map is represented using two colors: white cells indicate traversable locations, while black cells represent obstacles that the agent cannot enter. Traversable cells are connected to their valid neighbors, with movement allowed only in the four cardinal directions (north, east, south, and west), provided the neighboring cell is also traversable.

Together, these environments provide a controlled framework for investigating how spatial representations, value-guided plasticity, and contextual modulation interact to support navigation in biologically inspired spiking neural networks. The following section describes the neuromimetic spatial representations that provide state inputs to the learning mechanism.

### Neuromimetic Spatial Representations

Grid cells in the medial entorhinal cortex provide a periodic population code for spatial location, characterized by hexagonally arranged firing fields at multiple spatial scales (**Fig. 3**) (Fiete et al., 2008; Hafting et al., 2005; Moser et al., 2008; Solstad et al., 2006). Although individual grid cells are spatially ambiguous due to their repeating receptive fields, ensembles of grid cells with different spatial phases, scales, and orientations can uniquely encode location through combinatorial population activity (Leutgeb et al., 2005; Mathis et al., 2012; Rolls et al., 2006). Such distributed representations provide a compact and noise-tolerant encoding of spatial state and are widely believed to contribute to navigation and spatial memory(Aery Jones & Giocomo, 2023; Gorchetchnikov & Grossberg, 2007; Pouget et al., 2003; Zhang et al., 1998).

The present model adopts this principle by constructing a heterogeneous population of grid-cell-like inputs with multiple spatial scales, rotations, and offsets. These inputs provide a distributed spatial representation that serves as the primary state signal for downstream learning mechanisms.

Hippocampal place cells are thought to arise through integration of convergent spatial inputs from entorhinal cortex and related structures (Fenton et al., 2008; Knierim et al., 1995; McNaughton et al., 2006; O’Keefe & Dostrovsky, 1971; Solstad et al., 2006).

Place cells exhibit sparse spatial tuning, allowing individual neurons to respond selectively at specific locations within an environment. Inspired by this transformation from distributed entorhinal representations to localized hippocampal responses, the model incorporates an Association Cell (AC) population that integrates grid-cell activity through sparse fixed connectivity. The resulting AC activity forms more spatially selective internal representations that improve discrimination between locations and provide a suitable substrate for reinforcement learning.

### Maze Environments

Maze navigation provides a useful framework for studying biologically inspired reinforcement learning because it combines structured spatial representations with sparse and delayed reward signals(Anwar et al., 2026; Earl et al., 2026). From a computational perspective, successful navigation requires addressing three challenges: (1) distinguishing between spatial locations, (2) learning effective action sequences despite sparse feedback, and (3) supporting multiple navigation objectives within a shared environment.

To evaluate these challenges, we designed two complementary maze environments (**Fig. 1**). Both are discrete grid worlds containing obstacles that constrain movement and require multi-step planning to reach a goal location.

**Maze Type 1** (**Fig. 1A**) contains multiple starting locations and a single goal location. This environment primarily evaluates spatial state discrimination and learning under sparse reward conditions. The maze contains constrained pathways but no loops, allowing the effects of ΔQ-modulated learning to be examined in isolation.

**Maze Type 2** (**Fig. 1B**) contains multiple start-goal pairs within a shared spatial layout. In this setting, identical spatial locations may occur across several navigation objectives while requiring different actions depending on the active goal. This environment therefore evaluates the network’s ability to support multiple navigation policies within a shared architecture.

The proposed framework addresses these challenges using a common network structure. Maze Type 1 relies on grid-cell-derived spatial representations and ΔQ-modulated Hebbian plasticity, while Maze Type 2 introduces an additional Context Cell population that provides task-dependent modulation of internal representations without altering the underlying learning mechanism.

## Methods

### SNN + Q-Table Model

The proposed framework consists of a feedforward spiking neuronal network (SNN) coupled to a goal-conditioned Q-table (**Fig. 2**). The network contains four neuronal populations: Grid Cells (GCs), Association Cells (ACs), Context Cells (CCs), and Motor Cells (MCs). For Maze Type 1, the CC population is inactive and the model reduces to a three-population architecture (GC → AC → MC). For Maze Type 2, Context Cells provide task-dependent modulation that enables learning of multiple navigation objectives within a shared network.

**Figure 2.**
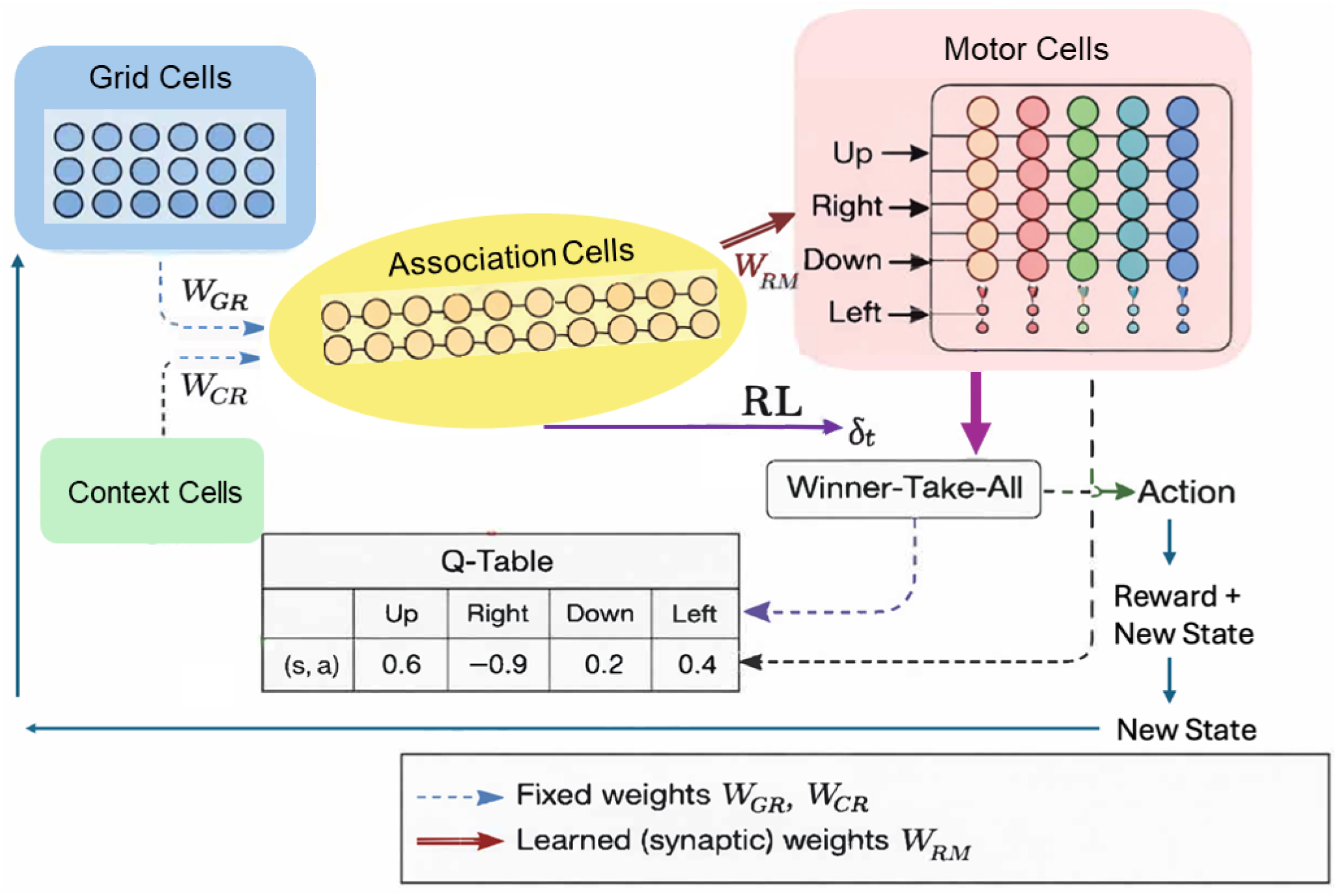
Schematic of the spiking neuronal network (SNN) architecture and reinforcement learning framework. Grid Cell (GC) inputs provide spatial representations of the agent’s current location, while Context Cells (CCs) provide task-dependent signals for multi-goal navigation. Both populations project through fixed synaptic connections (W_GR_, W_CR_) to an Association Cell (AC) population that generates distributed spatial representations. AC neurons project to Motor Cell (MC) populations through plastic synapses (W_RM_), which are modified by ΔQ-modulated Hebbian plasticity. Motor population activity is converted into an action through a winner-take-all mechanism. The selected action updates the agent’s state in the maze environment, producing a reward and new state that are used to update the Q-table. Changes in Q-values (ΔQ) provide the reinforcement signal that modulates synaptic plasticity in the AC→MC pathway.

**Figure 3.**
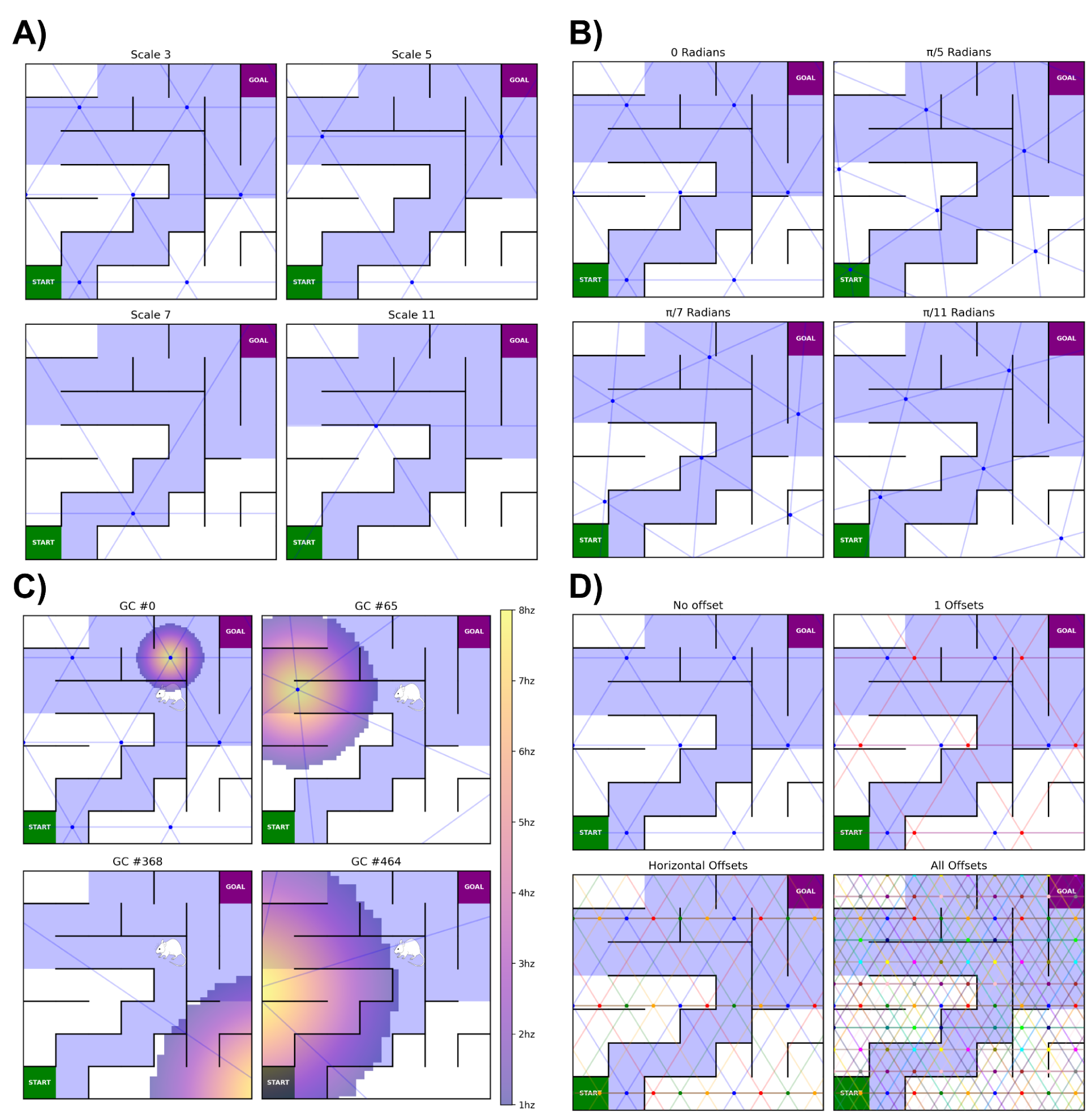
Grid-cell parameterization used to generate distributed spatial representations. Grid-cell modules differ in scale, rotation, offset, and firing-field location, producing heterogeneous population activity patterns that uniquely encode spatial position. Dots indicate firing-field centers and connecting lines illustrate the underlying hexagonal lattice structure. **A)** Grid-cell modules with scales 3, 5, 7, and 11, demonstrating how scale controls the spacing between firing-field centers. **B)** Grid-cell modules with different lattice orientations, showing how rotation alters the angular alignment of firing fields. **C)** Example firing-rate maps for individual grid cells overlaid on the maze environment. Color indicates firing rate (1-8 Hz), with peak activity centered on grid-field locations. **D)** Effect of spatial offsets within a grid-cell module. All cells within the module share the same scale and rotation but differ in spatial phase (different offsets along the x and y axis), generating distinct firing-field locations. Increasing the number of offsets expands coverage of the environment and contributes to the combinatorial population code used for spatial state representation (top to bottom, left to right, 1, 2, 4, and 16 GCs are plotted).

The architecture was designed to address three fundamental challenges of navigation. The first challenge is generation of distinct spatial representations that allow different locations to be reliably discriminated. This function is supported by the GC and AC populations, which transform distributed grid-cell activity into more selective location-dependent representations. The second challenge is learning under sparse reward conditions, where successful behavior depends on associating local neural activity with delayed consequences. This function is achieved through a goal-conditioned Q-table and ΔQ-modulated Hebbian plasticity acting on AC→MC synapses. The third challenge is learning multiple navigation objectives within a shared environment. This function is supported by the CC population, which introduces context-dependent modulation of AC activity and allows identical spatial locations to support different navigation policies.

Together, these components provide a unified framework for spatial representation, reinforcement learning, and goal-conditioned navigation. The following sections describe the neuronal populations, connectivity patterns, and learning mechanisms in greater detail.

#### Grid Cell Population

The Grid Cell (GC) population provides the primary spatial representation used by the model. Two experimentally observed properties of grid cells were incorporated into the implementation:

1. Grid cells are organized into modules whose neurons share common spatial scales and orientations but differ in spatial phase (offset).
2. Grid-cell activity varies systematically as an animal moves through a firing field.

Accordingly, the GC population was organized into modules, each defined by a unique spatial scale, orientation (rotation), and set of spatial offsets (**Fig. 3A,B,D**). Firing rates were determined by the agent’s position relative to nearby firing-field centers (**Fig. 3C**). This formulation captures the spatial selectivity of grid cells while omitting more complex temporal phenomena such as theta modulation and phase precession (Hafting et al., 2008), which are discussed further in the Discussion section.

To maximize representational diversity, grid-cell modules were assigned spatial scales based on prime numbers (Stensola et al., 2012). Because firing fields with prime-number spacing overlap less frequently than fields with common factors, this arrangement increases the uniqueness of the resulting population code. Five grid-cell modules were used with scales of 3, 5, 7, 11, and 13 (**Table 2**). Scale 2 was excluded because it produced excessively dense firing-field coverage and elevated baseline activity. All scales were multiplied by a global scaling factor of 0.5 to ensure that multiple firing peaks from each module were represented within the maze environment.

**Table 1.**
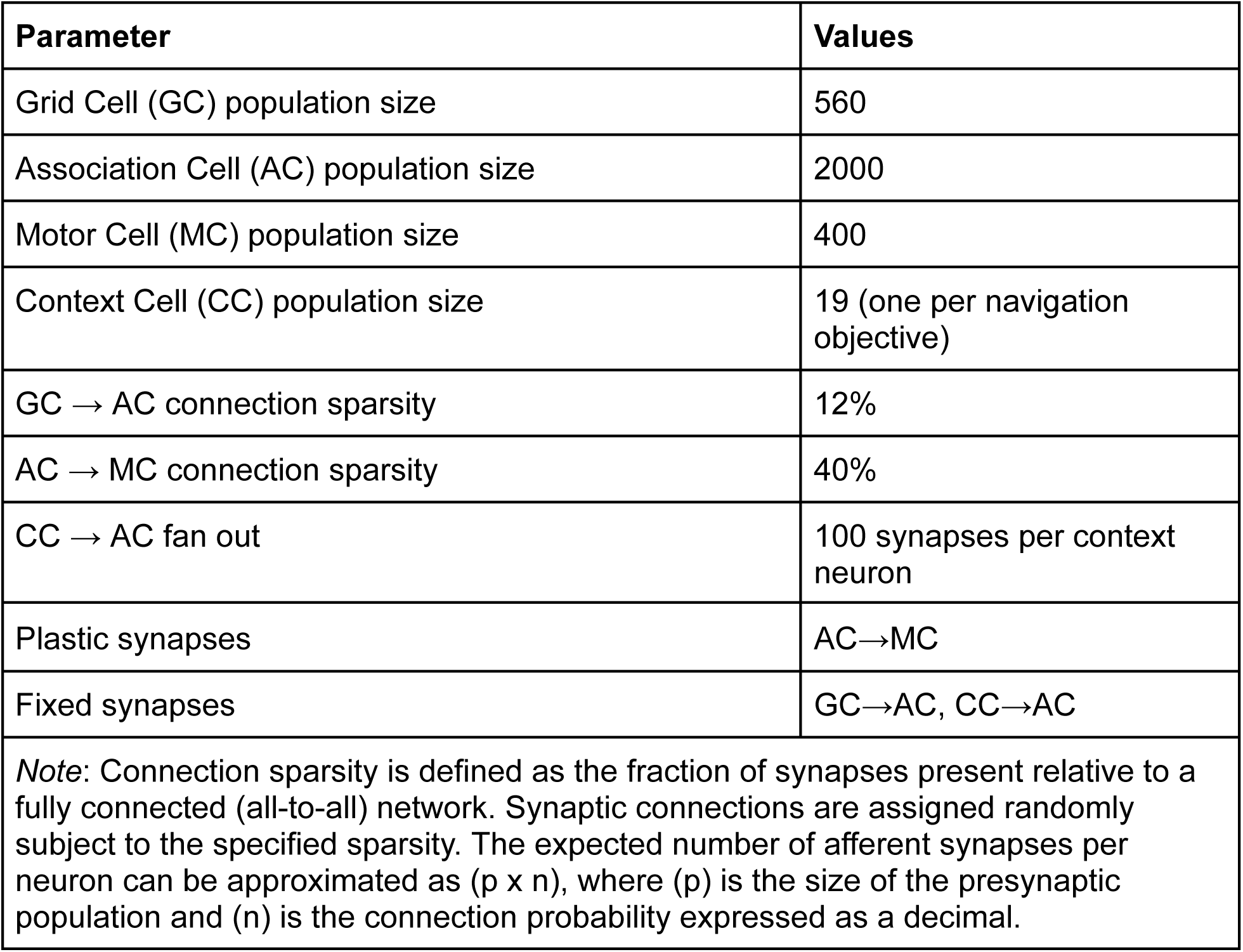
Population sizes and connection sparsity parameters for the SNN architecture.

| Parameter | Values |
| --- | --- |
| Grid Cell (GC) population size | 560 |
| Association Cell (AC) population size | 2000 |
| Motor Cell (MC) population size | 400 |
| Context Cell (CC) population size | 19 (one per navigation objective) |
| GC → AC connection sparsity | 12% |
| AC → MC connection sparsity | 40% |
| CC → AC fan out | 100 synapses per context neuron |
| Plastic synapses | AC→MC |
| Fixed synapses | GC→AC, CC→AC |
| *Note:* Connection sparsity is defined as the fraction of synapses present relative to a fully connected (all-to-all) network. Synaptic connections are assigned randomly subject to the specified sparsity. The expected number of afferent synapses per neuron can be approximated as $(p \times n)$, where $(p)$ is the size of the presynaptic population and $(n)$ is the connection probability expressed as a decimal. | |

**Table 2:** Parameters used to generate the Grid Cell (GC) population. Spatial scales define the spacing between firing-field centers, rotations determine lattice orientation, and offsets define spatial phase within each module. The global scale factor (g) is applied uniformly to all grid-cell scales, preserving their relative prime-number spacing while ensuring that multiple firing peaks are represented within the maze environment.

| Grid Cell Parameters |  |
| --- | --- |
| Parameter | Value |
| Spatial scales | 3, 5, 7, 11, 13 |
| Rotations | $0, \pi / 3, \pi / 5, \pi^*3 / 5, \pi / 7, \pi^*3 / 7, \pi / 11$ |
| Spatial offsets | 16 evenly spaced offsets arranged on a 4×4 grid |
| Field sharpness (k) | 1 |
| Global scale factor (g) | 0.5 |
| Total grid cells | 560 (5 scales x 7 rotations x 16 offsets) |

Firing fields were parameterized using two-dimensional Gaussian probability density functions (PDFs) centered at each firing-field peak (**Fig. 3D**) (Kropff & Treves, 2008). For a given spatial location, firing rates were determined by evaluating the Gaussian PDF associated with the nearest firing-field center. To minimize overlap between neighboring firing fields, the covariance matrix (C) was defined as:

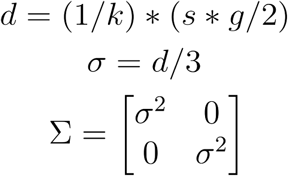

where *k* >= 1 controls the sharpness of the firing field, (s) is the grid-cell scale, and (g) is a global scaling parameter applied to all modules. The characteristic distance (d) was constrained to be no greater than half the spacing between adjacent firing peaks. The standard deviation was set to (σ = d/3), ensuring that approximately 99.7% of the Gaussian mass remained within the corresponding firing field. Values below 0.01 were rounded to zero when computing firing rates, further reducing overlap between neighboring fields.

Grid-cell firing rates were obtained by evaluating the Gaussian firing field associated with the nearest firing-field center at the agent’s current location. The resulting value was normalized to the range [0,1] and scaled by a maximum firing rate of 8 Hz. This value lies within the range of experimentally observed in-field firing rates for grid cells (Hafting et al., 2005). The firing rate for a grid cell at position (x,y) was therefore computed as: *f*(*x, y*) = (*p*(*x, y*)/*p_max_*) * *f_max_*, where *p*(*x, y*) denotes the value of the Gaussian firing field evaluated at the agent’s current location, and *p_max_* denotes the peak value of that firing field. Spike trains were generated by distributing *f*(*x, y*) spikes uniformly across a 1000 ms simulation interval.

#### Association Cell Population

The Association Cell (AC) population serves as an intermediate layer that transforms distributed grid-cell activity into more spatially selective representations. AC neurons receive sparse, fixed projections from the Grid Cell (GC) population, giving each AC a unique and unchanging set of afferent inputs. As a result, different spatial locations activate different subsets of AC neurons depending on the underlying pattern of GC activity.

ACs are modeled as leaky integrate-and-fire (LIF) point neurons (Dayan & Abbott, 2001). Their membrane properties were chosen to favor coincident activation, causing AC neurons to respond preferentially when multiple afferent GCs fire within a short temporal window. This mechanism allows ACs to integrate distributed grid-cell activity and generate sparse, location-selective responses analogous to hippocampal place-cell representations. The selectivity of these responses is primarily governed by the membrane potential decay constant and firing threshold parameters (**Table 3**).

**Table 3.** Leaky integrate-and-fire (LIF) neuron parameters(Hazan et al., 2018) used for Association Cells (ACs) and Motor Cells (MCs). Parameters were selected to promote sparse, coincidence-driven activation of AC neurons and competitive action selection within the MC population (MC neurons were assigned a zero refractory period to allow rapid competition between motor populations during action selection).

| Association Cell and Motor Cell LIF Parameters |  |  |
| --- | --- | --- |
| Parameter | AC | MC |
| Refractory period (ms) | 1 | 0 |
| Resting voltage (mV) | -64 | -64 |
| Reset voltage (mV) | -70 | -64 |
| Firing threshold (mV) | -45 | -49 |
| Membrane decay constant (ms) | 20 | 20 |
| Neurons per population | 2000 | 400 |

Through this transformation, the AC population converts distributed spatial codes into representations that are more readily associated with navigation decisions through downstream reinforcement learning.

#### Context Cell Population

To support learning across multiple navigation objectives within a shared environment, the model includes a Context Cell (CC) population that provides task-dependent input to the Association Cell (AC) layer (**Fig. 2**). Each navigation objective corresponds to a unique start-goal pair and is associated with a single context neuron.

Context neurons project to the AC population through fixed synaptic connections *W_CT_* (**Fig. 2**) with a connection probability of 0.05. Given 19 context neurons and 2000 AC neurons, this configuration produces approximately 100 outgoing synapses per context neuron (1900 total context-to-association synapses). These connections are not modified during learning and serve exclusively as a contextual input signal.

Contextual information is represented using a dedicated population of N_PATHS_ neurons, where each neuron corresponds to a unique navigation objective. For a given path (P), the context input is represented as a binary spike-train matrix of size (SIM_TIME * N_PATHS_), in which only the neuron associated with the active path is permitted to spike while all other context neurons remain silent.

The active context neuron emits spikes periodically at fixed intervals (every 20 simulation time steps; simulation time step = 1 ms), producing a regular spike train of approximately 50 Hz throughout the navigation episode. Although all navigation objectives share the same temporal spiking structure, they differ in the identity of the active context neuron. Context spike trains are delivered through a contextual input population and projected to the AC layer through fixed random synaptic connections.

The CC population does not encode spatial position, goal coordinates, or motor actions directly. Instead, it provides a low-dimensional contextual signal that interacts with the grid-cell-derived spatial representation within the AC population. Because identical spatial locations may occur across multiple navigation objectives, this contextual input allows AC activity to depend on both the agent’s location and the currently active navigation objective.

Synaptic plasticity occurs only in AC→MC connections and is modulated by the ΔQ teaching signal derived from the goal-conditioned Q-table. Consequently, Context Cells influence learning indirectly through their effect on AC population activity. This mechanism allows a shared AC population, motor population, and Q-table to support multiple navigation policies without requiring separate networks for each navigation objective.

#### Motor Cell Population

The Motor Cell (MC) population serves as the action-selection layer of the network. It is divided into four subpopulations corresponding to the four possible navigation actions: Up, Down, Left, and Right. Each motor subpopulation contains 100 neurons and is modeled using leaky integrate-and-fire (LIF) dynamics with parameters similar to those used for the AC population (**Table 3**).

Like AC neurons, MC neurons are configured to respond preferentially to coincident input from multiple upstream neurons. Consequently, the activity of each motor population reflects the combined influence of the currently active AC neurons and the learned AC→MC synaptic weights. Through ΔQ-modulated Hebbian plasticity, these synaptic connections become specialized so that different spatial representations preferentially activate the motor population associated with the appropriate navigation action.

Motor population activity is converted into a discrete behavioral action using a winner-take-all (WTA) mechanism. At each decision step, the motor subpopulation with the highest aggregate spiking activity is selected, and the corresponding action is executed in the environment. The WTA mechanism operates downstream of neural activity and is not itself subject to learning. Instead, it serves as a readout mechanism that transforms distributed population activity into one of the four discrete navigation actions.

### Reinforcement Learning

#### Learning Under Sparse Reward Conditions

Many reinforcement learning tasks provide dense feedback that can be used to directly reinforce or suppress synaptic activity. Maze navigation presents a more challenging setting because environmental feedback is sparse: most actions produce either no reward or a penalty, while positive reward is received only upon reaching the goal.

Under these conditions, local synaptic activity must be associated with outcomes that may occur many steps in the future.

To address this problem, the model incorporates a goal-conditioned Q-table derived from the Q-learning algorithm (Christopher J C & Dayan, 1992; Watkins & Dayan, 1992). The Q-table stores value estimates for state-action pairs, where larger values indicate actions expected to produce more favorable future outcomes. After each action, Q-values are updated using the Bellman equation (Bellman, 1957; Bellman & Kalaba, 1957, 1959), allowing information about future rewards to propagate backward through the state space. As a result, the Q-table provides value estimates that account for long-term consequences rather than immediate reward alone.

The resulting Q-values are not used directly to determine synaptic changes. Instead, changes in Q-values (ΔQ) are used to modulate Hebbian plasticity in the AC→MC pathway, allowing local neural activity to be associated with delayed navigation outcomes. The specific plasticity mechanism is described in the following section.

To evaluate learning under sparse reward conditions, Maze Type 1 was trained to navigate to a single goal location using a five-phase protocol (**Fig. 4**):

1. Exploratory navigation from starting location 1
2. Learning from starting location 1
3. Exploratory navigation from starting location 2
4. Learning from starting location 2
5. Recall and evaluation from starting location 1

**Figure 4.**
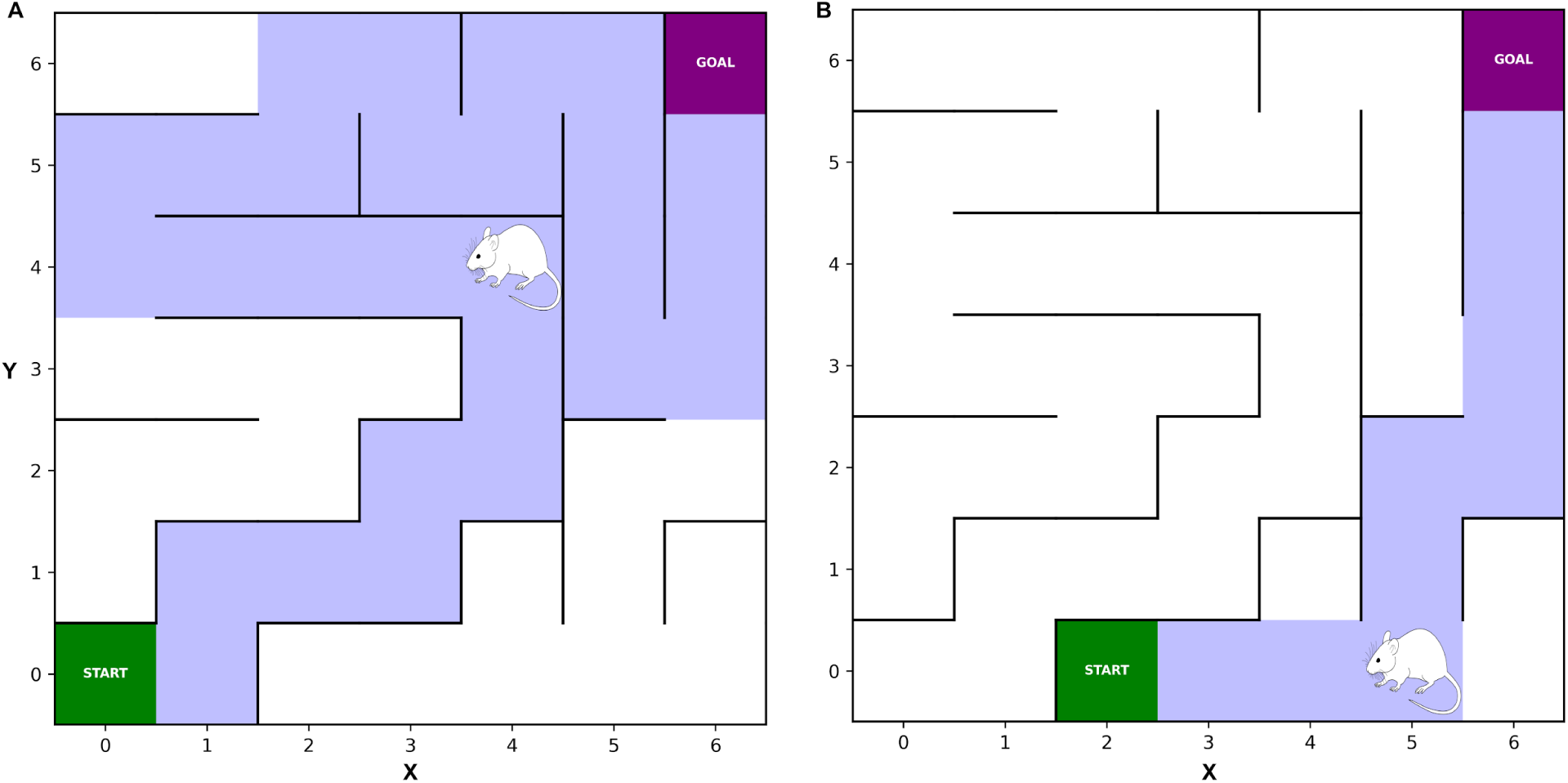
Two navigation tasks (A,B) used in Maze Type 1. The agent must learn to reach a common goal location (purple) from two distinct starting locations (green) within the same maze. Blue shading indicates the optimal path associated with each start location. These tasks are used to evaluate whether the network can learn and retain multiple navigation strategies within a shared spatial environment while minimizing interference between previously learned policies.

The exploratory phases (1 and 3) are intended to mimic initial exploration of an unfamiliar environment. During these phases, AC→MC plasticity is disabled and actions are selected randomly. Although behavior is not guided by the Q-table, Q-values continue to be updated based on state transitions and rewards encountered during exploration. These phases allow the agent to discover the maze structure and establish initial value estimates before synaptic learning is enabled.

During learning phases (2 and 4), ΔQ-modulated Hebbian plasticity is enabled.

Q-values continue to be updated, and changes in Q-values are used to modulate synaptic plasticity, allowing the network to associate local neural activity with actions that improve expected future reward.

Phase 5 evaluates retention of previously learned navigation behavior. After learning from the second starting location, the agent is returned to the original starting location and required to navigate using the previously acquired policy. This phase provides a measure of the extent to which learning one navigation path interferes with previously learned behavior and therefore serves as a test of memory retention within the shared network architecture.

#### ΔQ-Modulated Hebbian Plasticity

Learning in the network occurs through Hebbian plasticity applied to AC→MC synapses following each navigation decision (during active learning phases). Rather than using environmental reward directly, synaptic updates are modulated by changes in Q-values (ΔQ) obtained from the goal-conditioned Q-table. This approach allows local synaptic plasticity to incorporate information about long-term navigation outcomes.

The intuition behind this formulation is straightforward. A positive ΔQ indicates that the selected action moved the agent toward a state with higher expected future reward, whereas a negative ΔQ indicates movement toward a state with lower expected future reward. Consequently, positive ΔQ values reinforce the neural activity patterns that produced the selected action, while negative ΔQ values weaken those associations.

When ΔQ = 0, the transition provides little or no information regarding the relative quality of the selected action.

At each decision step, the grid-cell population is simulated according to the agent’s current location for a 1000 ms interval. The resulting GC activity drives the AC and MC populations, and the motor action is selected using the winner-take-all mechanism described above. Following execution of the selected action and the subsequent Q-table update, ΔQ is computed and used to modulate plasticity in AC→MC synapses.

To promote competition between motor populations, different plasticity rules are applied to synapses projecting to the selected motor population and to synapses projecting to non-selected motor populations. For synapses connecting presynaptic neuron (i) to postsynaptic neuron (j) within the selected motor population, synaptic weight updates are given by Δ*w_i,j_* = *x_i_* * *x_j_* * Δ*Q* * α, where *x_i_* and *x_j_* denote the number of spikes emitted by neurons *i* and *j*, respectively, and α is the learning rate between (0, 1]. For synapses projecting to all non-selected motor populations, synaptic weight updates are computed as Δ*w_i,j_* = *x_i_* * *x_j_* * Δ*Q* * α.

Under this learning rule, positive Δ*Q* values strengthen synapses contributing to the selected action while weakening competing action pathways. Conversely, negative Δ*Q* values weaken synapses contributing to the selected action and redistribute synaptic influence toward alternative actions. Through repeated interaction with the environment, this competitive process progressively associates spatial representations in the AC population with actions that improve expected future reward.

#### Goal-Conditioned Q-Table for Maze Type 2 Learning

The Maze Type 1 experiments use a spatially indexed Q-table because all training paths converge to a single goal location. In contrast, Maze Type 2 requires learning multiple navigation objectives within the same environment. Under these conditions, a Q-table indexed only by spatial location becomes insufficient, since identical locations may require different actions depending on the active goal. To address this limitation, the Q-table was extended to include both spatial position and goal identity in the state representation, enabling multiple goal-conditioned navigation policies to be learned within a single shared value structure.

The state was defined as *s* = (*g, p*), where (*g*) denotes the active goal location and (*p*) denotes the agent’s current spatial position. Q-values were therefore indexed as: *Q*(*g, p, a*), where (*a*) denotes one of the four navigation actions (Up, Down, Left, Right).

In practice, the Q-table was implemented as a dictionary mapping the tuple (*g, p*) to a vector of four action values. This goal-conditioned indexing allows the same spatial location to be associated with different action values depending on the active navigation objective. Consequently, distinct navigation policies can be learned without maintaining separate Q-tables for each task.

An additional advantage of this formulation is that navigation objectives sharing the same goal can reuse portions of the same value structure. Tasks with different starting locations but identical goals therefore contribute to a common set of value estimates, reducing redundancy while preserving goal-specific behavior.

Q-values were updated using the standard Q-learning update rule: *Q*(*s, a*) ← *Q*(*s, a*) + α(*r* + *γ max*(*a*′) *Q*(*s′, a′*) − *Q*(*s, a*)), where *s* is the current state, *a* is the action selected in the current state, *s*’ is the successor state reached after executing *a*, *a*′ indexes possible actions available from the successor state, *r* is the received reward, α is the Q-learning rate, and *γ* is the discount factor.

During learning, the Q-table provides value estimates that are used to compute (\Delta Q), which in turn modulates Hebbian plasticity in AC→MC synapses. Thus, the Q-table serves as a source of long-horizon value information while the network itself remains responsible for generating actions through MC population activity.

#### Runtime Methodology

At each decision step, the agent occupies a spatial location (x,y) within the maze.

Grid-cell spike trains are generated according to the spatial encoding procedure described in the Grid Cell Population section. These spike trains drive activity in the GC population, which propagates through the AC and MC populations via the network architecture shown in **Figure 2**.

Motor population activity is converted into a discrete action using the winner-take-all mechanism described above. The motor subpopulation producing the greatest aggregate activity determines the selected navigation action (Up, Down, Left, or Right). The agent is then moved to the corresponding location, the Q-table is updated, and synaptic plasticity is applied according to the ΔQ-modulated Hebbian learning rule (during active learning phases). This process is repeated until the goal is reached or the episode terminates.

#### Precomputation of GC and AC Activity

A naive implementation would regenerate GC spike trains and simulate AC activity each time the agent moves to a new location. However, the maze environments considered here are discrete rather than continuous, implying a finite set of possible spatial states.

To improve computational efficiency, GC spike trains were precomputed for every traversable location in the maze prior to training. Furthermore, because GC→AC synapses are fixed and AC neuron parameters remain unchanged throughout training, AC activity for a given spatial location is deterministic. Consequently, AC spike trains were also precomputed for all maze locations.

This precomputation substantially reduces simulation cost by avoiding repeated evaluation of identical spatial representations during training and testing.

#### Software and Computational Resources

The model was implemented using the BindsNET spiking neural network simulation framework (Hazan et al., 2018). Simulations were performed on a workstation equipped with an AMD Ryzen 9 7950X3D processor (16 cores, 32 threads, 4.2 GHz) and 32 GB of RAM. Under these conditions, a complete training session required approximately five minutes of runtime.

Source code, simulation scripts, and visualization tools are publicly available through GitHub and ModelDB (Hines et al., 2004).

## Results

We evaluated the proposed framework against the three navigation challenges introduced above: 1) generation of distinct spatial representations, 2) learning under sparse reward conditions, and 3) goal-conditioned navigation within a shared environment. Results are organized accordingly. First, we assess the spatial selectivity and separability of Grid Cell (GC) and Association Cell (AC) representations. Second, we examine the ability of ΔQ-modulated Hebbian plasticity to learn efficient navigation policies from sparse feedback. Finally, we evaluate whether Context Cell modulation and a goal-conditioned Q-table enable the network to learn multiple navigation objectives within a shared architecture. Together, these analyses assess the ability of the model to support flexible, goal-directed navigation across both maze environments.

### Goal 1: Spatial Selectivity of GC and AC Representations

The first navigation challenge requires the network to generate distinct neural representations for different spatial locations. While the quantitative analyses below are performed using Maze Type 1, the same Grid Cell (GC) and Association Cell (AC) architecture is used in Maze Type 2, where representational separability is equally important due to the larger number of navigation objectives.

To evaluate spatial selectivity, we simulated network activity at every location in the maze and identified active neurons, defined as cells that emitted at least four spikes during the simulation interval. We then quantified both the number of active neurons at each location and the overlap in active neuronal ensembles between pairs of locations. Effective spatial representations should satisfy two criteria: (1) sufficient neuronal activity to support downstream learning and action selection, and (2) low overlap between locations to enable reliable state discrimination.

Figure 5 illustrates the transformation from distributed grid-cell activity to sparse AC responses. At location (0,5), subsets of GCs exhibit temporally clustered firing due to overlapping grid-field activation. AC neurons integrate these convergent inputs and generate sparse, location-selective responses, producing activity patterns analogous to hippocampal place cells.

**Figure 5.**
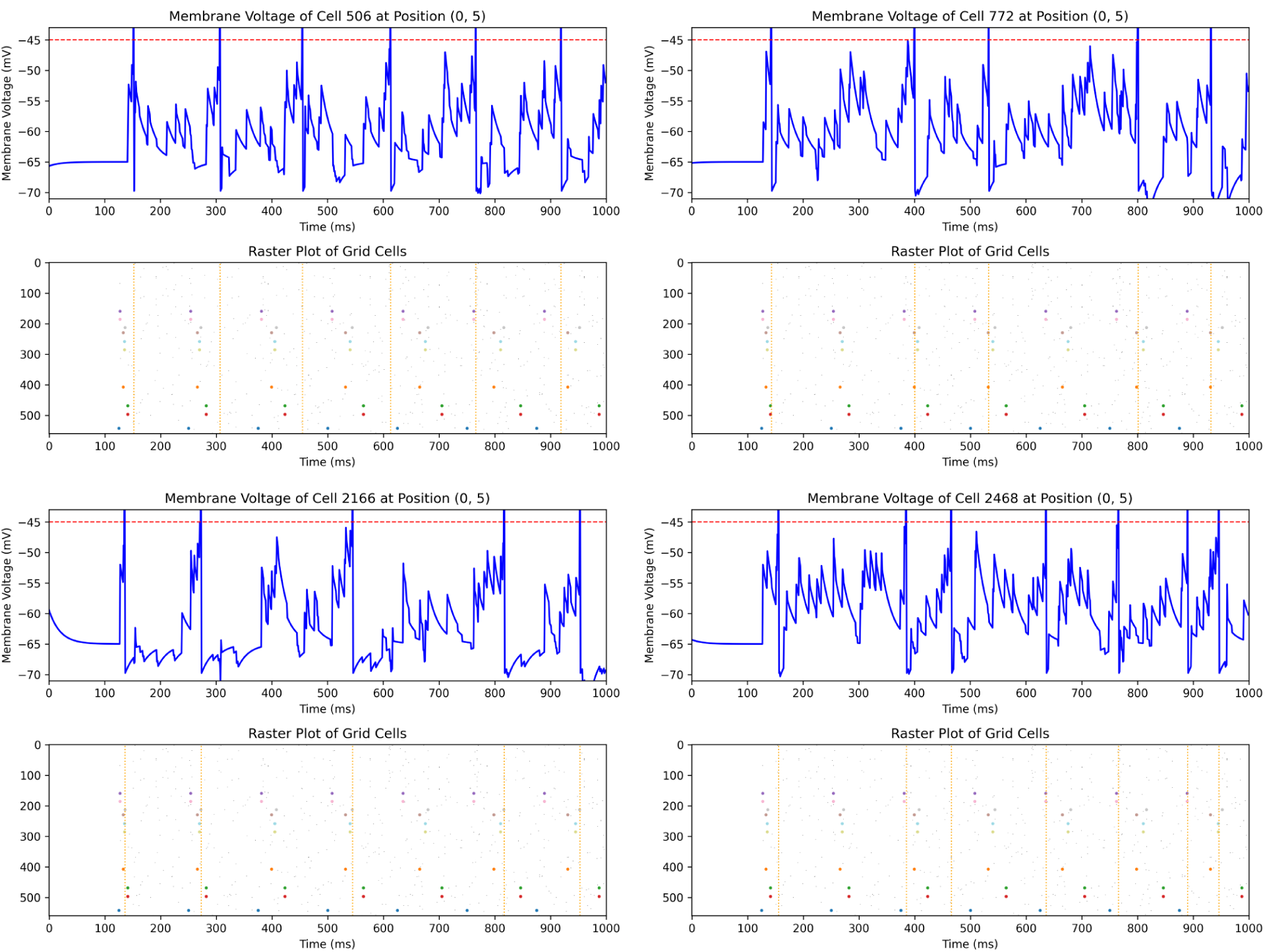
Emergence of place-cell-like responses in Association Cells (ACs) through integration of grid-cell activity. Membrane voltage traces of four active Association Cells (ACs) when the agent is positioned at coordinate (0,5). Blue curves show membrane voltage and the red dashed line indicates the firing threshold (-45 mV). Raster plots below each voltage trace show spikes from active Grid Cells (GCs), defined as cells that emitted at least four spikes during the simulation interval; inactive GCs are omitted for clarity. Colored dots indicate GC spike times, while vertical orange dashed lines denote AC spike times. At this spatial location, subsets of GCs exhibit temporally clustered firing due to overlapping grid-field activation. AC spikes occur primarily during periods of convergent GC activity, demonstrating that AC neurons integrate distributed grid-cell inputs to generate sparse, location-specific responses analogous to hippocampal place cells.

The resulting spatial representations are summarized in Figure 6 and Table 4. Figure 6A shows that both GC and AC populations maintain substantial activity across the maze, with an average of 52.3 active GCs and 58.3 active ACs per location. Thus, the AC layer preserves a rich representation of spatial state while transforming upstream grid-cell activity.

**Figure 6.**
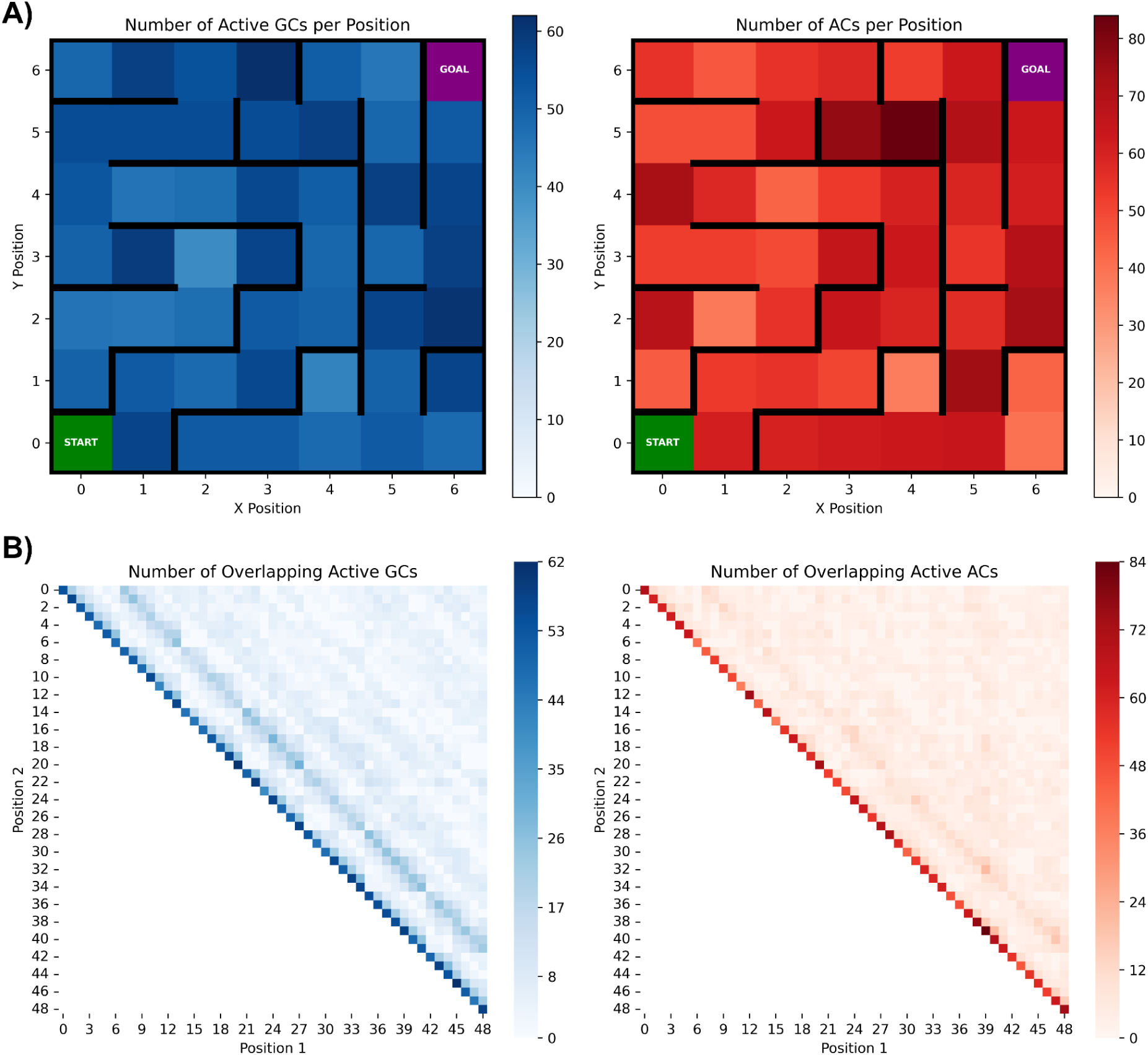
Spatial selectivity and representational separability of Grid Cell (GC) and Association Cell (AC) populations. **A)** Heat maps showing the number of active GCs (left, blue) and ACs (right, red) at each location in the 7x7 maze. A cell was considered active if it emitted at least four spikes during the simulation interval. Both populations maintain substantial activity throughout the environment, providing sufficient representation density for downstream learning. **B)** Pairwise overlap matrices showing the number of active GCs (left) and ACs (right) shared between all pairs of maze locations. Diagonal elements represent the total number of active cells at a given location, while off-diagonal elements represent overlap between distinct locations. The relatively low off-diagonal values indicate that different spatial positions are represented by distinct neuronal ensembles. Compared to the GC population, the AC population exhibits reduced overlap between locations while maintaining robust activity levels, demonstrating that the AC layer transforms distributed grid-cell inputs into more spatially selective, place-cell-like representations. These results support Goal 1 by showing that the network generates unique and discriminable neural representations across the maze environment.

**Table 4.** Numerical values and interpretations from **Fig 6**. *Encoding Quality Score* (bottom row) is calculated as the average number of active cells divided by the average number of overlapping active cells.

| Data | Grid Cells | Association Cells |
| --- | --- | --- |
| Min. Diagonal Value | 40 | 37 |
| Max Non-diagonal Value | 27 | 21 |
| Avg. Active Cells in all Positions | 52.3 | 58.3 |
| Avg. Overlapping Active Cells for all Position Pairs | 5.7 | 3.5 |
| Encoding Quality Score | 9.2 | 16.6 |

Figure 6B quantifies representational overlap between all pairs of maze locations. Diagonal elements represent the number of active neurons at a given location, whereas off-diagonal elements represent overlap between distinct locations. Both GC and AC populations exhibit substantially lower off-diagonal than diagonal values, indicating that different spatial positions are represented by distinct neuronal ensembles. The AC population further reduces overlap relative to the GC population, with average pairwise overlap decreasing from 5.7 active neurons for GCs to 3.5 for ACs (**Table 4**).

To summarize representational quality, we computed an Encoding Quality Score defined as the ratio of the average number of active neurons to the average pairwise overlap between locations. Higher values indicate representations that maintain strong activity while minimizing interference between spatial states. The Encoding Quality Score increased from 9.2 for the GC population to 16.6 for the AC population, demonstrating that the AC layer improves the separability of spatial representations generated by the upstream GC population.

Together, these results show that the network generates distributed yet distinct neural representations across the maze environment, providing a suitable substrate for downstream reinforcement learning.

### Goal 2: Long-Horizon Navigation Learning under Sparse Reward

To evaluate Challenge 2, we examined the ability of the network to learn efficient navigation policies using ΔQ-modulated Hebbian plasticity. Figure 7 shows the learning dynamics of a representative training session together with averages across 50 independently generated maze environments.

**Figure 7.**
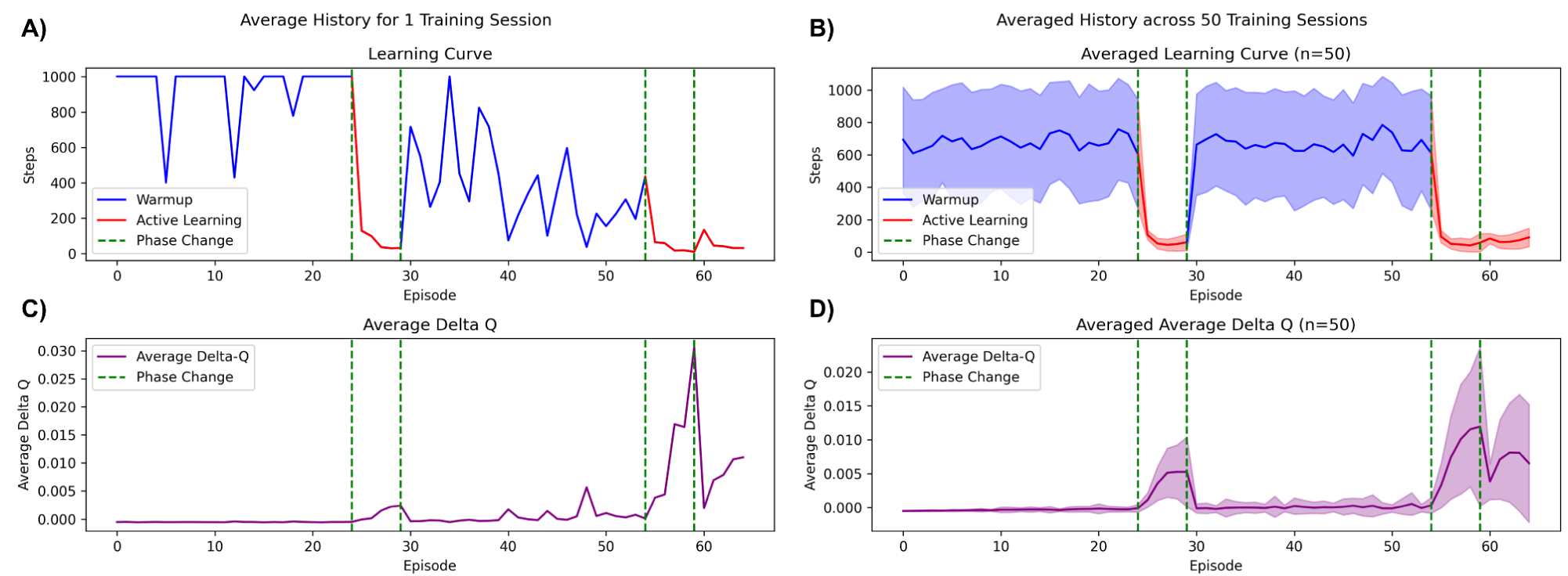
Learning performance and ΔQ dynamics during maze navigation. **A,B)** Learning curves showing the number of steps required to reach the goal as a function of training episode. **C,D)** Average ΔQ value per episode, computed by averaging ΔQ across all actions within an episode. Panels **A** and **C** show results from a single training session, while panels **B** and **D** show means across 50 independently generated maze environments; shaded regions indicate ±1 standard deviation. Blue curves denote exploratory warmup phases (Phases 1 and 3), during which synaptic plasticity is disabled and the Q-table is populated through random navigation. Red curves denote active learning phases (Phases 2, 4, and 5), during which ΔQ-modulated Hebbian plasticity is enabled. Green dashed lines indicate phase transitions, including changes in starting location. Across both individual and population-averaged results, active learning produces a rapid reduction in path length accompanied by positive ΔQ values, indicating that the network successfully associates spatial states with actions that increase expected future reward. The low variance observed during active learning demonstrates consistent acquisition of effective navigation policies across diverse maze environments.

The training protocol consisted of five phases. During **Phase 1**, the agent performed exploratory navigation from the first starting location. Synaptic plasticity was disabled, and only the Q-table was updated. Because reward information was initially unavailable and the AC→MC pathway had not yet been shaped by learning, behavior during this phase was effectively exploratory, resulting in substantial variability in the number of steps required to reach the goal across episodes. The purpose of this phase was to initialize the Q-table with value estimates that reflected the structure of the environment and could subsequently be used to generate ΔQ signals for synaptic learning.

During **Phase 2**, ΔQ-modulated Hebbian plasticity was enabled, allowing the network to learn a navigation policy from the first starting location (marked in red in Fig. 7). To prevent the agent from becoming trapped by imperfect Q-value estimates, action selection incorporated an ε-greedy exploration strategy. The exploration parameter was initialized to ε = 1 and decayed by a factor of 0.99 after every step, resulting in approximately a 60% probability of random exploration after 50 steps and approximately 10% after 230 steps. Throughout this phase, the Q-table continued to update while ΔQ values were used to modulate synaptic plasticity. This enabled the network to associate local neural activity with long-term navigation outcomes despite the absence of dense environmental feedback.

**Phases 3 and 4** repeated this procedure from a second starting location. Phase 3 consisted of exploratory warmup with Q-table updates only, while Phase 4 enabled ΔQ-modulated Hebbian learning for the second navigation path.

Finally, **Phase 5** evaluated memory retention after learning multiple navigation paths. The agent was returned to the original starting location and required to navigate using previously learned synaptic weights. No additional warmup phase was performed and synaptic plasticity between AC and MC populations was substantially reduced, limiting the network’s ability to rewrite previously learned connections. Consequently, this phase served as a direct test of whether learning the second path interfered with the first.

The agent successfully recalled the original navigation policy, demonstrating that previously acquired path information was retained. However, performance was modestly degraded. Excluding the first two episodes of each phase, average path lengths increased by 16.26 ± 4.70 steps (mean ± SEM) between Phase 2 and Phase 5. The first two episodes were excluded because Phase 2 necessarily begins before any path-specific synaptic optimization has occurred, resulting in transiently elevated path lengths. The observed increase in steps suggests partial interference between the two learned navigation policies and provides an estimate of the memory limitations of the current architecture. We revisit these interference effects in the Discussion section.

The resulting behavioral dynamics are shown in Figures 7 **and 8**. During the warmup phases (blue curves in Fig. 7), navigation remained inefficient and highly variable because actions were guided primarily by exploration and partially initialized Q-values. This behavior is also visible in Figure 8A, where the agent explores large portions of the maze without reaching the goal before the episode terminates at the maximum allowed 1000 steps. Once ΔQ-modulated Hebbian learning was enabled (red curves in Fig. 7), the number of steps required to reach the goal decreased rapidly. Figure 8B illustrates the resulting transition from exploratory behavior to efficient goal-directed navigation, with the agent following a substantially shorter path to the goal.

**Figure 8.**
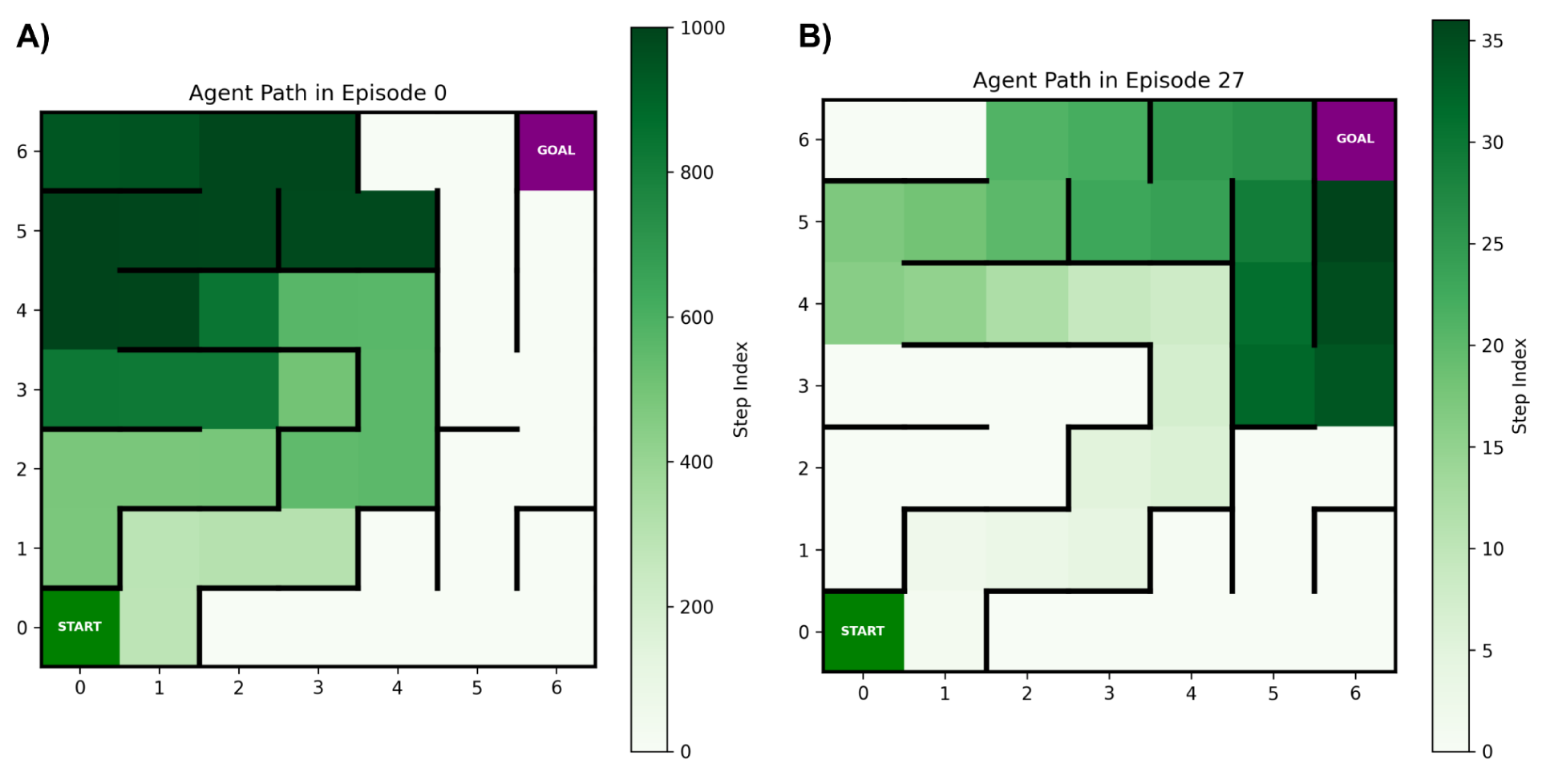
Transition from exploratory to goal-directed navigation. Heat maps showing the temporal progression of the agent’s trajectory during a representative warmup episode from Phase 1 **(A)** and an active learning navigation episode from Phase 2 **(B)**. Color indicates the order of visited locations, with lighter shades corresponding to earlier steps and darker shades to later steps in the episode. During the warmup phase **(A)**, the agent explores the maze through largely random movements, traversing many locations without reaching the goal and eventually terminating after the maximum allowed 1000 steps. Following two episodes of ΔQ-modulated Hebbian learning **(B)**, the agent reaches the goal through a substantially shorter and more direct trajectory. This figure illustrates the behavioral consequence of learning: a transition from inefficient exploratory search to efficient goal-directed navigation under sparse reward conditions.

The average ΔQ values shown in Figures 7C**,D** provide a complementary measure of learning progress. Because the Maze Type 1 environment contains no loops, positive ΔQ values generally correspond to actions that move the agent toward states with higher expected future reward, whereas zero or negative ΔQ values correspond to actions that do not improve progress toward the goal. Consequently, episodes with larger average ΔQ values tend to contain a greater proportion of productive navigation decisions and require fewer steps to reach the goal. Consistent with this interpretation, increases in average ΔQ occur primarily during active learning phases and coincide with substantial reductions in path length.

Figure 7 further demonstrates that these effects generalize across maze instances. Averaged results across 50 independently generated mazes (Fig. 7B**,D**) show substantial reductions in path length during active learning phases, accompanied by consistently positive ΔQ values. Variability is highest during warmup phases, when navigation behavior is dominated by exploration, and substantially lower during active learning. Together, these results indicate that ΔQ-modulated Hebbian plasticity reliably produces effective navigation policies across diverse maze environments.

The spatially selective representations identified in Figure 6 imply that different maze locations are associated with distinct subsets of active AC neurons. Because different locations may require different optimal actions, successful learning should produce location-specific changes in AC→MC connectivity that reflect the appropriate motor decision for each spatial state.

To examine this mechanism, Figure 9 shows the distribution of AC→MC synaptic weights at different stages of training for selected maze locations in the environment shown in Figure 1. Figure 9A corresponds to position **(4,4)**, where the optimal action is Left. Prior to learning, synaptic weights are approximately evenly distributed across motor populations. Following Phase 2, synaptic weight originating from active ACs becomes concentrated on the Left motor population, indicating that the network has associated the local AC representation with the correct navigation decision. Importantly, this weight specialization remains largely unchanged after Phase 4, despite subsequent learning of a second navigation path, indicating that the previously learned policy is retained.

**Figure 9.**
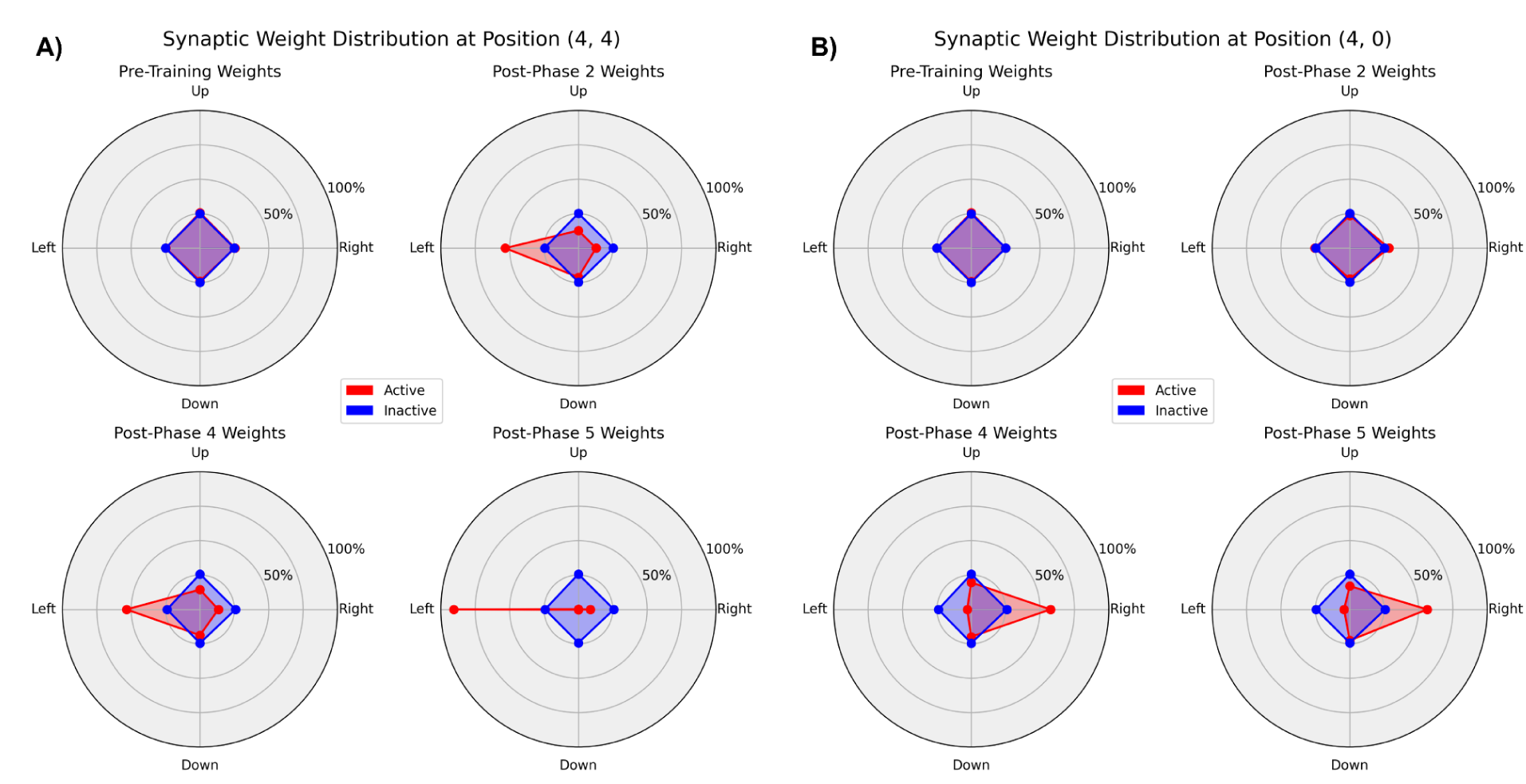
Location-specific reorganization and retention of AC→MC synaptic weights during learning. Radar plots show the distribution of synaptic weight assigned to the four motor populations (Up, Down, Left, Right) at different stages of training. The angular axis represents motor actions and the radial axis indicates the fraction of total AC→MC synaptic weight allocated to each motor population. Values were computed by summing all afferent AC→MC synaptic weights projecting to a motor population and normalizing by the total AC→MC synaptic weight. Red curves represent synapses originating from active Association Cells (ACs) at the specified location, while blue curves represent synapses from inactive ACs. **A)** Synaptic weight distributions at location (4,4), which lies on the first learned navigation path and for which the optimal action is Left. Following Phase 2 learning, weights from active ACs become concentrated on the Left motor population and remain largely preserved through Phases 4 and 5. **B)** Synaptic weight distributions at location (4,0), which lies on the second learned navigation path and for which the optimal action is Right. Consistent with its later introduction during training, strong weight specialization emerges only after Phase 4 and remains stable during Phase 5. Together, these results show that ΔQ-modulated Hebbian learning produces location-specific action preferences while preserving previously acquired policies after learning additional navigation paths.

Figure 9B shows the corresponding weight distributions at position **(4,0)**, where the optimal action is Right. Unlike position **(4,4)**, this location lies on the second learned path rather than the first. Consequently, strong weight specialization does not emerge until after Phase 4, when the second path has been learned. Once established, these weights remain stable during Phase 5, indicating that acquisition of the second navigation policy does not require overwriting previously learned AC→MC associations.

Together, these results provide a synaptic-level explanation for the behavioral improvements observed in Figures 7 and **8**. Distinct AC representations for different spatial locations (Fig. 6) are transformed through ΔQ-modulated Hebbian plasticity into location-specific action preferences (Fig. 9). The persistence of these synaptic weight distributions after learning multiple paths further suggests that the network can retain multiple navigation policies within a shared set of synaptic connections while limiting interference between previously acquired memories.

### Goal-Conditioned Navigation in Maze Type 2

Having demonstrated that the network generates distinct spatial representations (Challenge 1) and can learn efficient navigation policies under sparse reward conditions (Challenge 2), we next evaluate its ability to support multiple navigation objectives within a shared environment. To accomplish this, we activate the Context Cell (CC) population and evaluate the full four-population architecture in Maze Type 2.

Maze Type 2 consists of a two-dimensional environment containing obstacles and traversable regions, in which the agent must navigate from a specified start location to a designated goal location. Unlike Maze Type 1, which focuses on learning multiple paths to a single goal, Maze Type 2 requires learning multiple distinct start-goal pairs within the same spatial layout. Consequently, the same spatial location may occur in several navigation objectives while requiring different actions depending on the active goal. This environment therefore provides a test of whether Context Cell modulation and a goal-conditioned Q-table can support multiple navigation policies within a shared network architecture.

#### Multi-Path Learning in a Shared Environment

In Maze Type 2, each navigation objective is defined by a unique start–goal pair. Because many objectives share portions of the same environment, identical spatial locations can occur across multiple navigation tasks while requiring different actions depending on the active goal. Consequently, spatial representations alone are insufficient to uniquely determine the correct action, creating the potential for interference between learned policies.

To address this problem, the model augments the grid-cell-derived spatial representation with a task-dependent context signal. Context neurons project through fixed synaptic connections (W_cr_) to the Association Cell (AC) population, providing an additional source of input that identifies the currently active navigation objective. In the present implementation, each navigation objective is represented by a single context neuron. During a navigation episode, only the context neuron associated with the active objective emits spikes, producing a regular spike train of approximately 50 Hz. Although the context population is small relative to the 2000-neuron AC population, each active context neuron projects to 100 randomly selected AC neurons through sparse divergent connectivity. This allows a low-dimensional context signal to influence a distributed subset of the high-dimensional AC representation.

The resulting context signal does not encode spatial position directly; instead, it modulates AC activity by interacting with the underlying grid-cell representation. As a result, the same spatial location can produce different AC population activity patterns under different navigation objectives.

This context-dependent modulation allows the network to associate identical spatial states with different actions while maintaining a shared AC population, shared motor population, and shared goal-conditioned Q-table. The following analyses examine whether this mechanism produces distinguishable neural representations and supports learning of multiple navigation policies within the same environment.

#### Learning Performance and Generalization

The model demonstrated substantial improvements in navigation performance across most navigation objectives following active learning (Fig. 10). In Maze Type 2, each navigation objective corresponds to a unique start-goal pair and is referred to as a path (Paths 0-18). Final evaluation path lengths were generally low, indicating that the network successfully learned efficient goal-directed navigation policies. Most paths exhibited greater than 70% reduction in path length relative to warmup exploration, demonstrating effective optimization of behavior through ΔQ-modulated Hebbian plasticity. Across-seed variability remained low for many paths, indicating that learning performance was generally reproducible across random initializations. A small number of paths, particularly Paths 0 and 10, exhibited elevated variability across seeds, suggesting increased sensitivity to initialization or stronger interference arising from the underlying maze structure.

**Figure 10.**
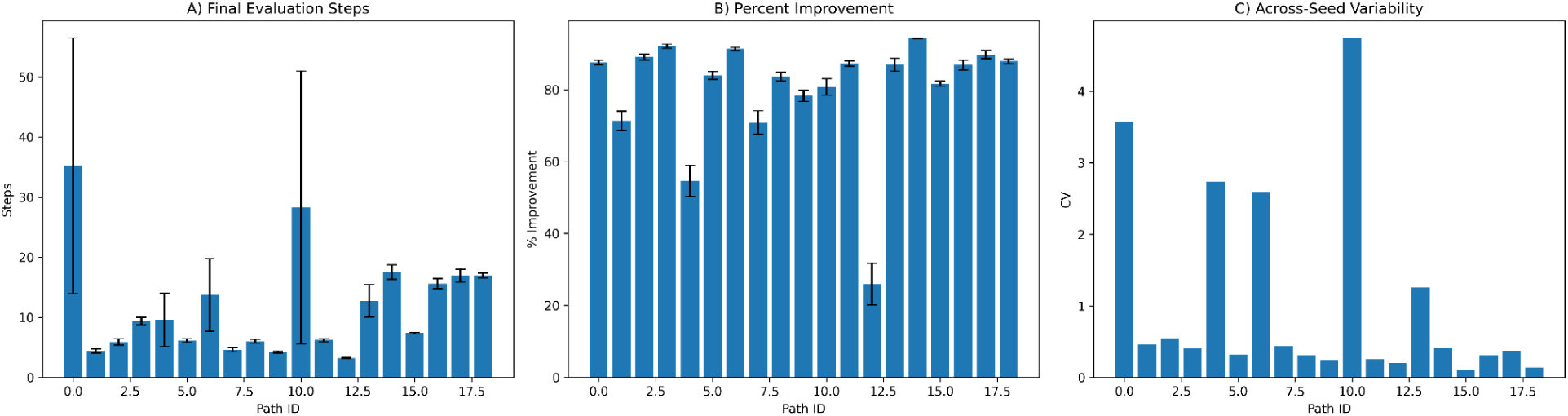
Multi-path navigation performance using a shared goal-conditioned Q-table. Results are shown for 19 navigation paths trained and evaluated across 35 random seeds in Maze Type 2. Each path corresponds to a distinct start-goal pair learned within the same environment using a shared Q-table and Context Cell modulation. **A)** Final evaluation performance, measured as the mean number of steps required to reach the goal after learning. Error bars indicate standard error of the mean (SEM) across seeds. Lower values correspond to more efficient navigation policies. **B)** Percent improvement relative to warmup exploration, quantifying the reduction in path length achieved through learning. Error bars indicate SEM across seeds. Higher values indicate greater learning-induced improvement. **C)** Across-seed variability, measured as the coefficient of variation (CV = standard deviation / mean) of evaluation-phase step counts. Lower values indicate more consistent performance across random initializations. Most navigation objectives exhibit substantial improvement and low final path lengths, demonstrating that the network can learn multiple goal-directed policies within a shared spatial environment while maintaining generally robust performance across seeds.

These results demonstrate that the network can learn multiple paths within a shared spatial environment using a common AC population, motor population, and goal-conditioned Q-table. Although all paths are learned within the same network, contextual modulation allows identical spatial locations to be associated with different actions when required by different start–goal pairs. Consequently, spatial structure can be reused across tasks without collapsing multiple policies into a single undifferentiated solution.

While Figure 10 summarizes final navigation performance across all paths, Figure 11 illustrates the learning dynamics for a representative path (Path 1; start = **(2,0)**, goal = **(2,3)**). **Figures 11A** and **11B** show the evolution of path length across episodes for a single training session and the corresponding average across 35 random seeds, respectively. During the warmup phase, navigation behavior is highly variable and inefficient, reflecting exploratory interactions with the environment prior to learning. Following the onset of active learning, path lengths decrease substantially, indicating the acquisition of more efficient navigation strategies. Similar improvements are observed in the across-seed average, demonstrating that learning dynamics are reproducible across random initializations. Evaluation performance remains stable following training, indicating successful retention of the learned navigation policy.

**Figure 11.**
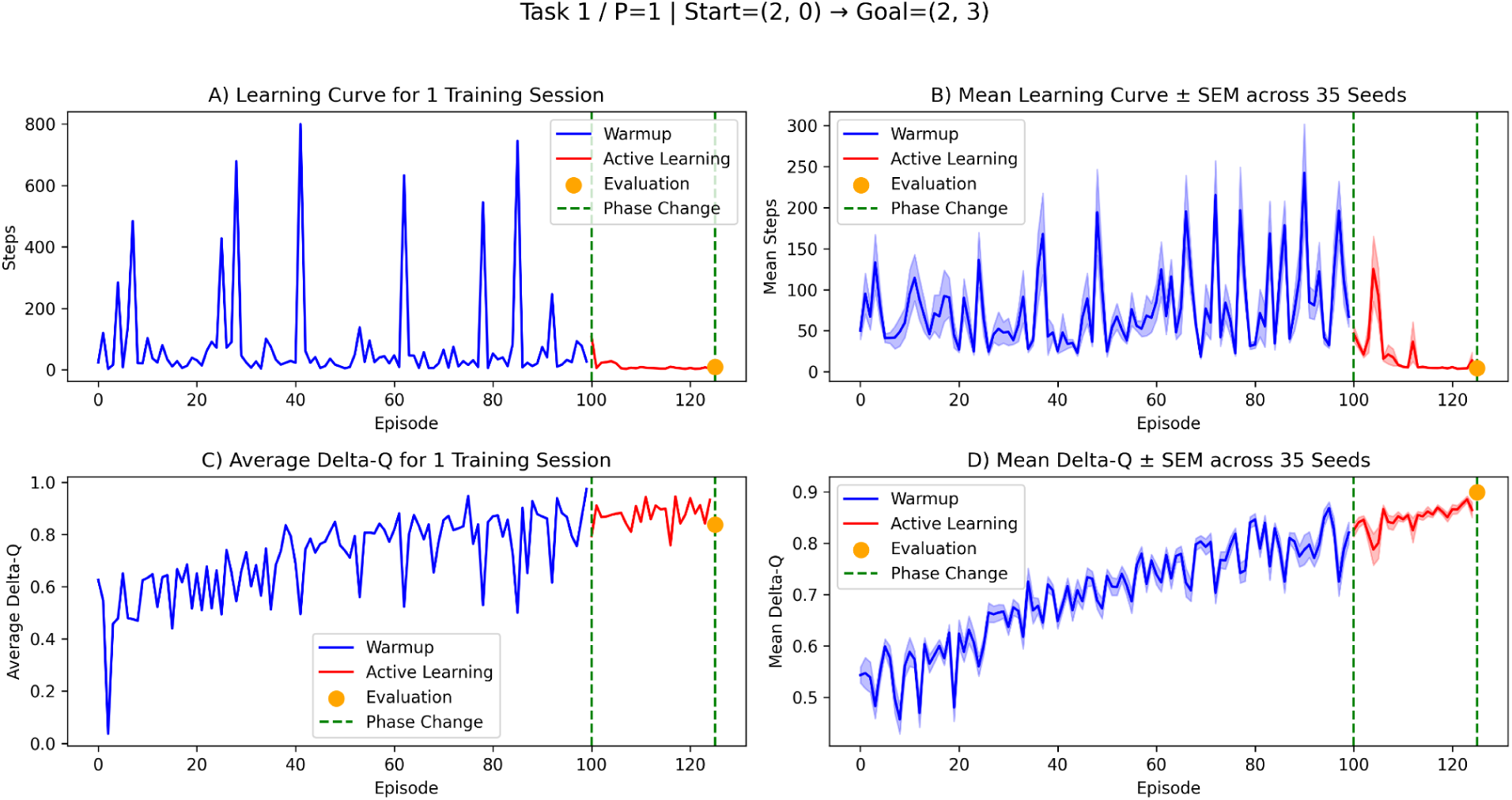
Learning dynamics and ΔQ evolution for a representative navigation task. Results are shown for Path 1 (start = **(2,0)**, goal = **(2,3)**) during Maze Type 2 training. **A)** Number of navigation steps required to reach the goal during a single training session. **B)** Mean learning curve across 35 random seeds, with shaded regions indicating ± SEM. Blue curves denote warmup exploration, red curves denote active learning, orange markers indicate evaluation performance, and green dashed lines indicate phase transitions. Following the onset of ΔQ-modulated Hebbian learning, path lengths decrease rapidly and remain low during evaluation, indicating successful acquisition of an efficient navigation policy. **C)** Average ΔQ values across episodes for a single training session. **D)** Mean ΔQ dynamics across 35 random seeds (± SEM). Increasing ΔQ values during training indicate progressive refinement of state-action value estimates and stronger alignment between selected actions and expected future reward. The close agreement between individual-session and population-averaged trajectories demonstrates that both the behavioral improvements and associated ΔQ dynamics are robust across random initializations. Together, these results illustrate how ΔQ-guided plasticity transforms exploratory behavior into stable goal-directed navigation.

Figures 11C and **11D** show the corresponding evolution of the average ΔQ signal for a single training session and the corresponding average across 35 random seeds. As training progresses, ΔQ values increase, reflecting a growing tendency for the agent to select actions associated with higher expected future reward. Following the onset of active learning, ΔQ values remain elevated and exhibit reduced variability, consistent with the formation of increasingly effective navigation policies. The close agreement between individual-session and population-averaged trajectories further indicates that the learning dynamics are robust across random initializations.

Together, Figures 10 and **11** demonstrate that the proposed architecture supports learning of multiple paths within a shared environment while maintaining reproducible learning dynamics across seeds. The accompanying increases in ΔQ and reductions in path length indicate that context-modulated spatial representations and ΔQ-guided plasticity jointly support efficient goal-directed navigation under sparse reward conditions.

#### Context-Dependent Neural Representations with a Shared Q-Table

Maze Type 2 was designed to determine whether multiple navigation objectives could be learned within a shared reinforcement learning framework. In this setting, all paths were learned using a single goal-conditioned Q-table. Because the state representation included both spatial location and goal identity, the same spatial coordinate could be associated with different action values depending on the active navigation objective. Consequently, multiple paths could share portions of the same environment while maintaining distinct navigation policies.

To examine how contextual information influences internal neural representations, we analyzed activity within the Association Cell (AC) population at a fixed spatial location, (2,0) (Fig. 12). **Figure 12A** shows the deviation in spike count from the across-path mean for the (**K=200**) most strongly modulated AC neurons across all trained paths. Rather than exhibiting identical activity across navigation objectives, subsets of AC neurons display systematic increases or decreases in firing depending on the active path. The resulting vertical band structure indicates that specific neurons consistently differentiate among navigation objectives, demonstrating context-dependent modulation of AC population activity.

**Figure 12.**
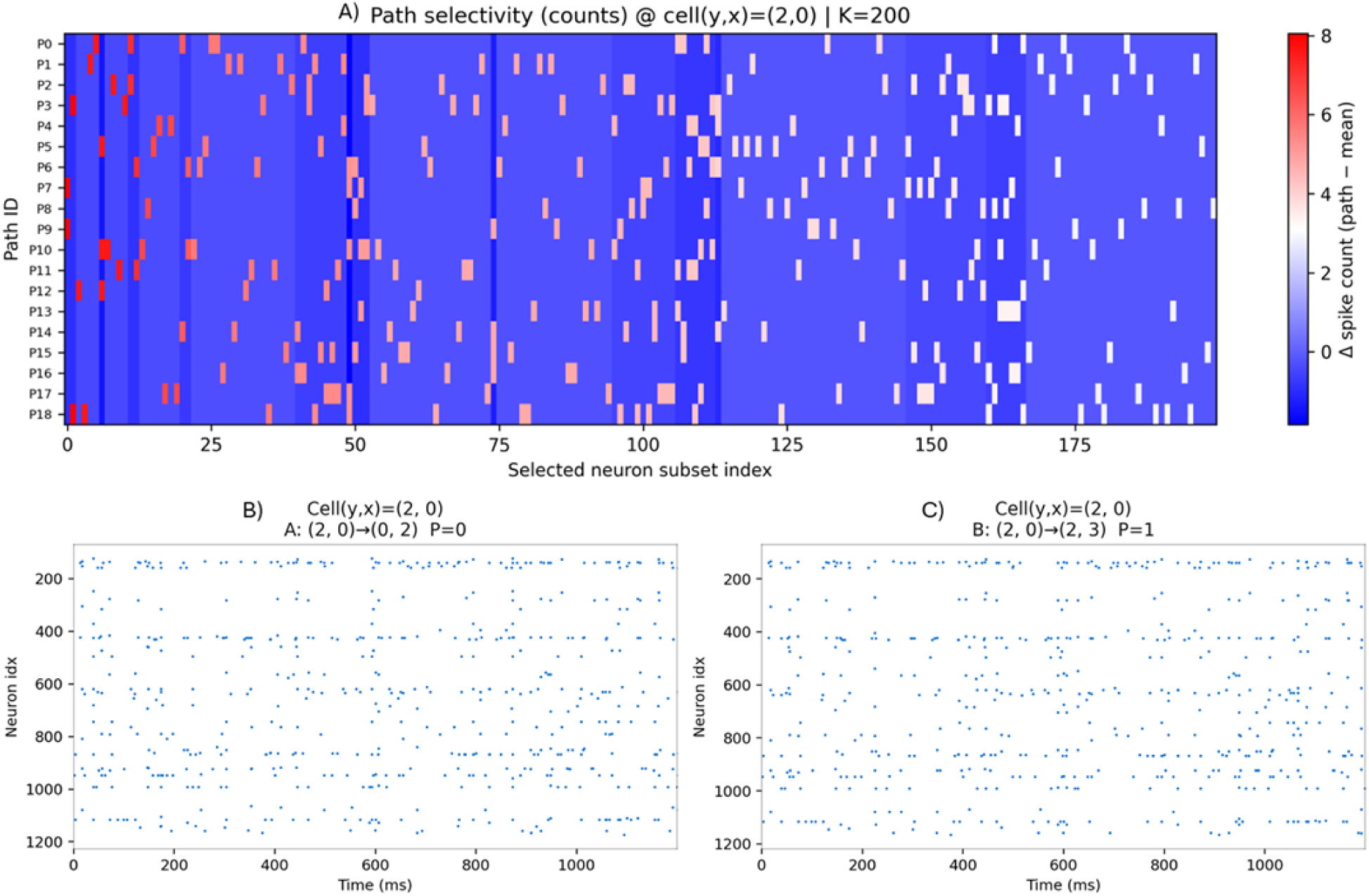
Context-dependent modulation of Association Cell (AC) activity at a shared spatial location. **(A)** Path-selective spike-count modulation for the **(K=200)** most strongly modulated AC neurons at spatial location **(2,0)**. Rows correspond to navigation paths (P0-P18) and columns correspond to selected neurons. Color indicates the deviation in spike count from the across-path mean (Δ spike count). Warmer colors indicate neurons that are more active than average for a given path, while cooler colors indicate reduced activity. Although all paths share the same spatial location, subsets of AC neurons exhibit systematic path-dependent modulation, producing distinguishable population activity patterns across navigation objectives. **(B,C)** AC spike rasters recorded at the same spatial location **(2,0)** under two different navigation contexts. Panel **B** corresponds to Path P0 **(2,0)→(0,2)**, and panel **C** corresponds to Path P1 **(2,0)→(2,3)**. Each dot represents a spike from an AC neuron (y-axis: neuron index; x-axis: time in ms). Despite identical spatial input, the resulting spatiotemporal activity patterns differ across paths, demonstrating that contextual input modulates AC population dynamics. Together, these results show that the same spatial state can generate distinct neural representations depending on the active navigation objective, enabling a shared network and goal-conditioned Q-table to support multiple navigation policies within the same environment.

Importantly, these path-dependent activity patterns emerge despite the use of a shared AC population, shared motor population, and shared goal-conditioned Q-table. The context signal therefore modifies how spatial information is represented within the AC layer, allowing the same spatial location to be associated with different downstream actions when required by different navigation objectives.

Figures 12B and **12C** provide complementary evidence by comparing AC spike rasters at the same spatial location, **(2,0)**, under two different navigation objectives. Although the spatial input is identical in both cases, the resulting spatiotemporal activity patterns differ between Path P0 (**(2,0)** → **(0,2)**) and Path P1 (**(2,0)** → **(2,3)**). These differences indicate that AC activity is influenced by both spatial input and contextual modulation rather than by spatial position alone.

To quantify the magnitude of this effect, we compared AC population activity between Paths P0 and P1 using cosine similarity of spike-count vectors. The resulting similarity value of 0.937 indicates that the two representations remain highly similar overall, reflecting their shared spatial input. At the same time, the residual differences are sufficient to distinguish navigation objectives and support different behavioral policies. Thus, contextual modulation does not completely remap the underlying spatial representation. Instead, it introduces subtle but consistent perturbations to a largely shared population code.

Together, these results provide a mechanistic explanation for the behavioral performance observed in Figures 10 and **11**. Context-dependent modulation of AC activity allows identical spatial locations to be represented differently under different navigation objectives, enabling a shared network and shared goal-conditioned Q-table to support multiple navigation policies while limiting interference between overlapping paths.

### Summary of Results

Taken together, the results from both maze environments demonstrate that the proposed framework addresses the three navigation challenges considered in this study. The GC and AC populations generate distinct spatial representations that support reliable state discrimination (Challenge 1), while ΔQ-modulated Hebbian plasticity enables learning of efficient navigation policies under sparse reward conditions (Challenge 2). In Maze Type 2, Context Cell modulation introduces task-dependent variations into AC population activity, allowing identical spatial locations to support different navigation objectives within a shared architecture (Challenge 3).

Across both environments, the model exhibits robust learning dynamics, efficient navigation performance, and stable retention of previously learned policies. These results suggest that distributed spatial representations, context-dependent modulation, and ΔQ-guided synaptic plasticity play complementary roles in supporting flexible goal-directed navigation within a biologically inspired spiking neuronal network.

## Discussion

We developed a biologically inspired spiking neuronal network (SNN) that combines grid-cell-based spatial representations, ΔQ-modulated Hebbian plasticity, and context-dependent modulation to address three core challenges of navigation: spatial state discrimination, learning under sparse reward conditions, and learning multiple navigation objectives within a shared environment. Across two complementary maze environments, the model generated distinct spatial representations, learned efficient navigation policies from delayed reinforcement signals, and successfully supported multiple goal-directed behaviors within a common network architecture.

The results suggest that these three components play complementary computational roles. First, the Association Cell (AC) population transformed distributed grid-cell activity into more spatially selective representations, increasing representational separability while maintaining robust activity levels across the environment. Second, ΔQ-modulated Hebbian plasticity enabled the network to associate local neural activity with long-term navigation outcomes despite sparse environmental feedback. Finally, Context Cell modulation introduced task-dependent variations into AC population activity, allowing identical spatial locations to support different navigation policies through subtle modifications of an otherwise shared spatial representation.

Although the model successfully addressed all three navigation challenges considered in this study, several important biological and computational limitations remain. Below we discuss four directions that may further improve both the biological realism and functional capabilities of the framework:

1. Improving grid-cell spike train realism
2. Replacing Hebbian plasticity with STDP-based learning
3. Introducing additional biologically motivated cell populations and circuit connectivity
4. Implementing the Q-table as a neuronal network

### Simplified Grid-Cell Spike Trains

A major simplification of the present model is the representation of grid-cell activity. In our implementation, grid-cell firing rates are determined by the agent’s distance from the nearest grid-field center and scaled between 0 and 8 Hz. Spikes are then distributed uniformly across a 1000 ms simulation window. This approach captures the spatial selectivity and periodic firing-field structure of grid cells but omits several temporal features observed in vivo (Hafting et al., 2005).

Experimental studies have shown that grid-cell activity is strongly modulated by hippocampal-entorhinal theta oscillations, producing temporally structured spike trains rather than uniformly distributed spikes (Hafting et al., 2008).In addition, both firing rate and spike timing vary systematically across a grid field, with spike timing exhibiting phase relationships to ongoing theta rhythms (Hasselmo et al., 2007). These temporal dynamics are thought to contribute to spatial coding, coordination of neuronal ensembles, and information transfer between entorhinal and hippocampal circuits (Cutsuridis et al., 2010).

The simplified spike trains used here were chosen to improve stability and reduce model complexity while isolating the contributions of spatial representations, ΔQ-modulated plasticity, and contextual modulation. Despite this simplification, the network generated distinct spatial representations (Fig. 6), learned efficient navigation policies under sparse reward conditions (**Figs. 7-9**), and supported multiple navigation objectives through context-dependent modulation of AC activity (Fig. 12). Nevertheless, because grid-cell activity provides the primary source of input to the network, the temporal structure of these inputs likely influences downstream neural dynamics, including the formation of place-cell-like representations and the operation of synaptic plasticity mechanisms.

Incorporating more biologically realistic theta-modulated grid-cell spike trains therefore represents an important direction for future work. Such inputs may alter the temporal organization of AC activity, improve coordination between neuronal ensembles, and provide a more appropriate substrate for temporally sensitive learning rules such as spike-timing-dependent plasticity (STDP)(Traub & Cunningham, 2026). More generally, introducing realistic temporal structure would allow future studies to examine how oscillatory dynamics contribute to the interaction between spatial representation, reinforcement learning, and memory formation in hippocampal-entorhinal circuits.

### STDP versus ΔQ-Modulated Hebbian Plasticity

In previous work, we successfully applied spike-timing-dependent reinforcement learning (STDP/RL) to tasks including simulated arm control, Cartpole, and Pong (Anwar et al., 2022; Haşegan et al., 2022; Neymotin et al., 2013). In those environments, reward signals varied continuously with behavioral progress, providing dense temporal feedback that could be used to reinforce specific patterns of pre- and post-synaptic activity. Under such conditions, temporally precise plasticity rules were effective at associating neural activity with successful outcomes.

Maze navigation presents a substantially different learning regime. Reward signals are sparse, often occurring only after extended sequences of actions, and successful behavior depends on associating local neural activity with delayed consequences. In the present model, grid-cell inputs were represented using simplified spike trains that lacked the temporal structure typically observed in hippocampal-entorhinal circuits. As a result, the precise spike-timing relationships required by STDP were not consistently present in the network activity. Under these conditions, we found that ΔQ-modulated Hebbian plasticity provided a more stable learning mechanism because synaptic updates depended on correlations between pre- and post-synaptic activity rather than the exact timing of individual spikes.

Importantly, this choice should not be interpreted as evidence that STDP is unsuitable for navigation learning. Rather, it reflects a limitation of the current model’s temporal dynamics. Experimental studies suggest that theta oscillations organize activity throughout hippocampal-entorhinal circuits and may provide the temporal structure necessary for spike-timing-dependent learning (Buzsáki, 2002, 2010; de Almeida et al., 2009; O’Keefe & Recce, 1993). Such oscillations can coordinate neuronal ensembles by constraining spikes to specific phases of the theta cycle, potentially creating the timing relationships required for effective STDP.

A natural next step is therefore to combine more biologically realistic theta-modulated grid-cell inputs with STDP-based reinforcement learning. Introducing oscillatory structure into the network could improve temporal coordination between neuronal populations, stabilize activity propagation through the circuit, and provide a more suitable substrate for temporally precise plasticity mechanisms. Determining whether STDP can match or exceed the performance of ΔQ-modulated Hebbian plasticity under these conditions remains an important direction for future work.

### Missing Biological Components and their Potential Benefits

The present model intentionally abstracts many features of the medial entorhinal cortex (MEC) and hippocampus in order to isolate the interactions between spatial representations, contextual modulation, and ΔQ-guided learning. While this simplification facilitated analysis of the learning mechanism, it omits much of the cellular and circuit diversity present in the biological system.

In vivo, the MEC, dentate gyrus (DG), CA3, and CA1 contain multiple classes of excitatory and inhibitory neurons whose interactions contribute to circuit stability, pattern separation, oscillatory dynamics, and information routing (Klausberger & Somogyi, 2008; Petilla Interneuron Nomenclature Group et al., 2008). By contrast, the current model contains only simplified excitatory populations and therefore lacks many of the regulatory mechanisms present in biological hippocampal-entorhinal circuits.

One consequence of this simplification is that the model relies primarily on sparse connectivity and cellular thresholds to produce distinct spatial representations. Although this approach was sufficient to generate location-selective AC activity (Fig. 6) and context-dependent modulation of shared representations (Fig. 12), inhibitory circuitry could potentially improve both representational precision and learning stability. For example, inhibitory interneurons are known to contribute to competition between neuronal ensembles, sharpen population responses, and regulate activity propagation throughout hippocampal networks. Similar mechanisms could help reduce representational overlap, improve separation between navigation objectives, and enhance robustness as the environment size and number of learned paths increase.

More broadly, the MEC, DG, and CA3 contain a substantially richer diversity of cell types and microcircuit motifs than represented here (Romani et al., 2024; Schneider et al., 2012). These populations support functions including pattern separation, pattern completion, gain control, and oscillatory coordination, all of which may influence spatial learning and memory(Buzsáki, 2002; McNaughton et al., 2006; Mizuseki & Buzsaki, 2009; Rolls, 2016). Incorporating additional biological detail therefore represents more than an effort to improve realism. Such mechanisms may provide computational advantages that become increasingly important as navigation tasks grow in complexity and scale.

### Q-Tables as Neuronal Networks

A central component of the present framework is the use of a goal-conditioned Q-table to provide value estimates that guide synaptic plasticity. Rather than learning directly from sparse environmental rewards, the network receives ΔQ signals derived from changes in Q-values, allowing local Hebbian plasticity to incorporate information about long-term navigation outcomes. This mechanism proved sufficient to support learning across both maze environments and enabled the network to associate distributed spatial representations with effective navigation policies.

The Q-table itself, however, remains an explicitly engineered component that exists outside the neural network. While this abstraction provides a practical mechanism for estimating future reward, it does not address how similar computations might be implemented within biological circuits. The basal ganglia has long been implicated in action selection, reward-based learning, and reinforcement signaling (Schultz et al., 1997), and receives dopaminergic inputs that modulate plasticity in ways broadly consistent with value-based learning mechanisms (Frank et al., 2004; O’Reilly & Frank, 2006; Paul Sands et al., 2023). Although the present model does not attempt to reproduce basal ganglia circuitry, the functional role of the Q-table is analogous in that both systems provide information about expected future outcomes that can influence behavior and learning.

A natural next step is therefore to replace the tabular Q-learning component with a neural implementation capable of generating similar value signals. Recent work has demonstrated Q-learning-like computations in spiking neuronal systems (Shin et al., 2026), suggesting a possible path toward a fully neural implementation of the framework. Such an extension would allow future studies to investigate how value learning, spatial representations, and synaptic plasticity interact within a unified neural architecture and may provide a more biologically grounded account of hippocampal-basal ganglia interactions during navigation.

### Future Directions

Several limitations of the current model suggest promising directions for future work. One important question concerns memory capacity and interference between learned navigation policies. The Phase 5 results revealed a measurable degradation in performance when recalling the first learned path after training on a second path, with average path lengths increasing from approximately 20 steps during initial learning to 30-40 steps during recall. This finding suggests that learning multiple navigation policies within a shared set of synaptic weights introduces a degree of interference, consistent with the broader challenge of catastrophic forgetting in neural systems (Parisi et al., 2019). Importantly, the model did not exhibit complete forgetting. Previously learned paths remained navigable, and the synaptic analyses in Figure 9 demonstrated that many location-specific AC→MC weight specializations were retained following acquisition of an additional path. Nevertheless, the observed performance degradation indicates that learning one navigation policy partially alters synaptic resources used by another, placing practical limits on the memory capacity of the current architecture.

This issue becomes particularly relevant in Maze Type 2, where up to 19 paths are learned within a shared network. Context Cell modulation partially mitigates interference by introducing task-dependent perturbations to Association Cell activity, allowing overlapping spatial states to support distinct policies. However, the degree to which interference scales with the number of learned paths remains unknown. Future work should systematically characterize memory capacity as a function of path count, spatial overlap, context dimensionality, and Association Cell population size. Such analyses would help determine whether the current architecture scales gracefully as the number of navigation objectives increases or whether additional mechanisms are required to preserve previously acquired policies.

Beyond memory capacity, several biologically motivated extensions may improve both the realism and computational capabilities of the model. Incorporating inhibitory interneurons and oscillatory dynamics may improve representational precision, stabilize network activity, and provide the temporal structure required for STDP-based learning. Likewise, replacing the engineered Q-table with a neural value-learning circuit would reduce reliance on non-neural components while enabling investigation of biologically grounded reinforcement mechanisms.

More broadly, the present work reflects a complementary modeling philosophy to many existing hippocampal and entorhinal cortex models. Numerous biophysical models of the trisynaptic circuit have focused on reproducing cellular properties, network connectivity, and emergent electrical dynamics with increasing biological realism (Dura-Bernal et al., 2024; Gandolfi et al., 2023). Such models have provided important insights into circuit organization and neural computation. In contrast, the current study begins with a simplified architecture designed to achieve functional learning on navigation tasks, and then uses the resulting behavior as a foundation for identifying which biological mechanisms may be necessary to reproduce or extend that functionality.

The future directions discussed above follow this philosophy. Incorporating inhibitory interneurons and oscillatory dynamics (Bezaire et al., 2016; Neymotin et al., 2011), introducing more realistic temporal structure into grid-cell activity, and replacing the engineered Q-table with a neural implementation of value learning (Shin et al., 2026) would each increase biological realism while preserving the functional benchmarks established here. This bidirectional approach from function toward biology, rather than only from biology toward function, may help clarify which cellular and circuit mechanisms are essential for flexible spatial learning and which primarily refine or stabilize existing computations.

The present results demonstrate that a relatively compact SNN, combining distributed spatial representations, context-dependent modulation, and ΔQ-guided plasticity, is sufficient to address several core challenges of navigation. Future work will determine the extent to which these capabilities persist as additional biological detail is introduced, and whether more realistic circuit architectures can preserve the functional advantages demonstrated here while providing deeper insight into the neural mechanisms underlying spatial learning and memory.

## Conclusion

We presented a biologically inspired spiking neuronal network model that combines grid-cell-derived spatial representations, ΔQ-modulated Hebbian plasticity, and context-dependent modulation to address three fundamental challenges of navigation: spatial state discrimination, learning under sparse reward conditions, and learning multiple navigation objectives within a shared environment. Across two complementary maze environments, the model generated distinct spatial representations, learned efficient navigation policies from delayed reinforcement signals, and successfully supported multiple goal-directed behaviors using a shared network architecture.

The results suggest that these components play complementary computational roles. The Association Cell population transformed distributed grid-cell activity into more separable spatial representations, ΔQ-guided plasticity associated those representations with long-term navigation outcomes, and Context Cell modulation enabled identical spatial locations to support different navigation policies. Together, these mechanisms allowed the network to learn and retain multiple navigation objectives despite sparse environmental feedback and overlapping trajectories.

More broadly, this work demonstrates that relatively simple spiking neural networks can combine biologically motivated representations with value-guided learning mechanisms to solve challenging navigation tasks. By focusing first on functional performance and then progressively reintroducing biological detail, this framework provides a foundation for investigating how hippocampal-entorhinal circuits, reinforcement learning mechanisms, and context-dependent neural dynamics interact to support flexible spatial behavior.

## Acknowledgements

Research supported by ARL Cooperative Agreement W911NF-22-2-0139, ARL/ORAU Fellowship, NIH R01DC012947, R01DC019979, R01MH134118-01, NIH R01NS128924-01, and P50MH109429.

## Notes

### Competing Interest Statement

The authors have declared no competing interest.

